# RNA surveillance controls 3D genome structure via stable cohesin-chromosome interaction

**DOI:** 10.1101/2021.05.31.446489

**Authors:** Yujin Chun, Sungwook Han, Taemook Kim, Yoonjung Choi, Daeyoup Lee

## Abstract

The 3D architecture that the genome is folded into is regulated by CTCF, which determines domain borders, and cohesin, which generates interactions within domains. However, organisms lacking CTCF have been reported to still have cohesin-mediated 3D structures with strong borders. How this can be achieved and precisely regulated are yet unknown. Using *in situ* Hi-C, we found that 3’-end RNA processing factors coupled with proper transcription termination are a cis-acting determinant that regulates the localization and dynamics of cohesin on the chromosome arms. Loss of RNA processing factors, including nuclear exosome and Pfs2, destabilizes cohesin from the 3’-ends of convergent genes and results in decreased cohesin-mediated domain boundaries. We observed the co-localization between Rad21 and a wide range of 3’end RNA processing/termination factors. Further, deletion of Rrp6 leads to cohesin redistribution to facultative heterochromatin, resulting in improper domain boundaries. Importantly, we observed that knockdown of Rrp6^Exosc10^ caused a defect in cohesin binding and loss of local topologically associating domains (TADs) interactions in mouse embryonic stem cells. Based on these findings, we propose a novel function of the RNA surveillance pathway in 3D genome organization via cohesin complex, which provides the molecular basis underlying the dynamics of cohesin function.

## Introduction

The mammalian genome is spatially organized into complex three-dimensional (3D) architecture encompassing compartments, topologically associating domains (TADs), and chromatin loops that allow for proper gene regulation (Dixon et al., 2012; Lieberman-Aiden et al., 2009; Nora et al., 2012; Rao et al., 2014). The 3D architecture is characterized by the convergent binding of the CCCTC-binding factor (CTCF) at the boundaries of chromatin domains and by the action of cohesin (Nora et al., 2017; Parelho et al., 2008; Rao et al., 2014; Rubio et al., 2008; Schwarzer et al., 2017; Wutz et al., 2017). The recent loop extrusion model suggested that cohesin extrudes loops until blocked by convergently oriented CTCF, which pre-marks the TAD borders (Fudenberg et al., 2016; Goloborodko et al., 2016; Sanborn et al., 2015). Cohesin and CTCF thereby maintain stable anchoring of chromatin loops and form discrete interaction domains.

Interestingly, these TAD-like structures and chromatin loops are also found in organisms that lack CTCF homologs. From *S.cerevisiae* to *S.pombe*, *C.elegans*, and *A. thaliana*, these organisms reveal discrete self-interacting domains with varying domain lengths independent of CTCF (Costantino et al., 2020; Hsieh et al., 2015; Rowley et al., 2017; Rowley and Corces, 2016). The cohesin, a ring-shaped protein complex, mediates the stable cohesion of sister chromatids and loop extrusion by holding distant DNA fragments in close spatial proximity (Murayama et al., 2018; Onn et al., 2008). In fission yeast, cohesin peaks are preferentially enriched at 3’-ends of convergent genes, and cohesin mediates the formation of interaction domains, termed ‘globules’, to organize chromatin architecture during interphase (Mizuguchi et al., 2014; Tanizawa et al., 2017). Even in Drosophila where CTCF homolog exists, Drosophila genome lacks evidence for the the colocalization of CTCF and cohesin complex as well as CTCF-mediated loop extrusion mechanism (Bartkuhn et al., 2009; Matthews and White, 2019). These findings imply that the formation and maintenance of chromatin interaction domains would require CTCF-independent regulatory factors (e.g., cohesin) or relevant biological processes, such as transcription/chromatin states, RNAPII or other transcription complex.

However, it is not fully resolved how cohesin determines the 3D chromatin organization and what mediates the stable cohesin localization in eukaryotes that lack CTCF or of which the CTCF does not play a major role. The cohesin accumulation at the 3’-end of convergent genes is achieved by passive translocation of cohesin along with RNAPII (Gullerova and Proudfoot, 2008; Schmidt et al., 2009). One possibility is the active role of transcription processes (including initiation, elongation, and termination) in regulating 3D chromatin structure and positioning of cohesin (Bhardwaj et al., 2016). Recent study suggested the importance of proper positioning of cohesin and RNAPII via transcription elongation in the formation of gene loops (Rowley et al., 2019). In addition, defects in transcription elongation cause mislocalization of cohesin and disruption of 3D chromatin interactions (Heinz et al., 2018). However, it has been suggested that not only the transcription elongation process which translocates cohesin together with elongating RNAPII, but also the 3’-end RNA processing machinery coupled with transcription termination plays a role in controlling genome topology and genome stability. In yeast, transcription terminator and CPF 3’-end processing machinery were reported to mediate gene looping (O’Sullivan et al., 2004; Tan-Wong et al., 2012). Given that loss of cohesin complex across the chromosomes results in missegregation of chromosomes and lagging chromosomes (Hadjur et al., 2009; Haering et al., 2008; Losada et al., 1998; Peters et al., 2008), involvement of various 3’-end processing and termination related factors in chromosome missegregation (Graham et al., 2009; Kinoshita et al., 1991; Mukarami et al., 2007; Ohkura et al., 1989, 1988; Shobuike et al., 2001; Wang et al., 2005; Win et al., 2006) imply an intimate relationship with cohesin.

Rrp6 and Dis3, which together form the nuclear exosome complex (Liu et al., 2006), are known to be involved in various RNA-related processes, including degradation of irregularly terminated transcripts (Vasiljeva and Buratowski, 2006) and noncoding cryptic unstable transcripts (CUTs) (Colin et al., 2011), and transcription termination of variety of noncoding RNAs (Fox et al., 2015). Along with RNA processing and termination-related functions, they also function in chromosome segregation (Graham et al., 2009; Mukarami et al., 2007). Pfs2, which encodes the *S. pombe* homolog of the WDR33 in mammalian CPSF complex, recognizes AAUAAA polyadenylation site (PAS) together with CPSF30 (Chan et al., 2014; Clerici et al., 2018; Schönemann et al., 2014). Defects in the CPSF complex lead to defects in transcription termination by ultimately causing extensive transcriptional readthrough genome-wide (Eaton et al., 2020). The loss of Pfs2 reportedly results in chromosome missegregation in *S. pombe* (Wang et al., 2005), indirectly implying a relationship between cohesin and beyond transcription elongation event. Thus, we focused our attention on elucidating the role of 3’-end mRNA processing linked with proper transcription termination at cohesin dynamics and 3D chromatin structure.

Here, we show that RNA processing factors involved in transcription termination contribute to the CTCF-independent regulation of cohesin localization and dynamics. Importantly, 3’-end RNA processing factors and proper transcription termination enables highly coordinated 3D genome architecture by regulating cohesin-mediated interaction domain formation. We confirmed that this relationship is also conserved in mouse embryonic stem cells. Finally, we provide a new molecular basis for control the dynamics of cohesin biology through uncovering a relationship between mRNA surveillance and cohesin complex.

## Result

### 3’-end RNA processing factors regulate the localization of the cohesin complex at the 3’-ends of convergent genes in euchromatin

To explore the relationship between 3’-end mRNA processing and chromatin localization of cohesin, we performed chromatin immunoprecipitation sequencing (ChIP-seq) for FLAG-tagged Rad21 (the kleisin subunit of cohesin complex) using mutant yeast with defects in the nuclear exosome complex (Rrp6 and Dis3) and the 3’-end formation of mRNA (Pfs2). Given that Dis3 and Pfs2 are essential and to avoid introducing bias from cell-cycle arrest, we grew the *dis3-54* and *pfs2-11* mutant cells at their semi-permissive temperatures (30°C and 32°C, respectively). Rad21 peaks identified in wild type cells showed a dominant (∼45%) distribution and strong enrichment at the 3’-ends of convergent genes and are regularly positioned across the genome as previously reported (Schmidt et al., 2009) (Figure 1 – figure supplement 1A). For further analysis, we mainly focused on the Rad21 sites (peaks) at intergenic sites where genes with opposite strands converge (converging intergenic sites). We detected a significant decrease in Rad21 intensity at the 3’-ends of convergent genes in *rrp6*Δ and *pfs2- 11* (67% and 58% fold decrease in *rrp6*Δ and *pfs2-11* cells, respectively), compared to those in wt cells (Figure 1A-C). In *dis3-54* mutant cells, reduction of Rad21 was relatively weaker than *rrp6Δ* due to a weak semi-permissive mutant of Dis3 (Figure 1A-C). Thus, we additionally employed a tetracycline-regulated system (Zilio et al., 2012) to modulate *dis3^+^* expression and check by conventional ChIP-qPCR whether the temporal depletion of Dis3 affects Rad21 occupancy. Consistent with previous report (Lemay et al., 2014), depletion of Dis3 by over ∼80% caused the severe defect in transcription termination, resulting in readthrough transcripts (Figure 1 – figure supplement 1B-C). Also, we observed the two-fold reduction of Rad21 enrichment upon Dis3 knockdown (Figure 1 – figure supplement 1D). We additionally performed ChIP-seq of FLAG-tagged Psc3^SCC3/STAG1/STAG2^ (an accessory subunit of the cohesin complex) and found that Psc3 occupancy in the *rrp6Δ* mutant was significantly reduced (Figure 1 – figure supplement 1E-F). These findings imply that 3’-end RNA processing factors are involved in regulating the occupancy of the cohesin complex at converging intergenic sites.

**Figure 1.**
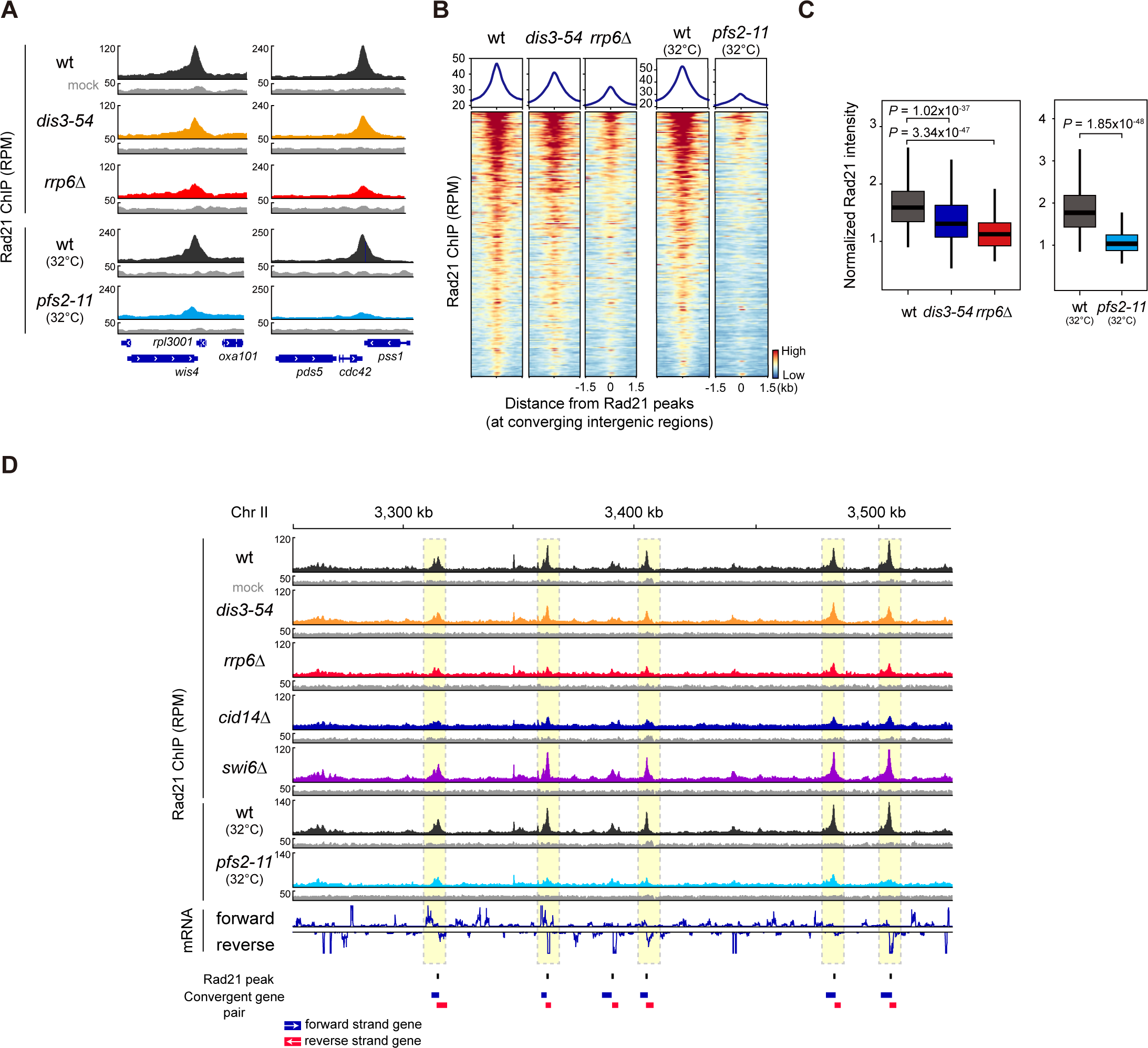
3’-end RNA processing factors regulate the localization of the cohesin complex at the 3’- ends of convergent genes in euchromatin. **(A)** Rad21 ChIP analysis at the 3’-end of convergent genes (*wis4*-*rpl3001* and *cdc42*-*pss1*) in *rrp6*Δ, *dis3-54* (30°C), and *pfs2-11* (32°C) compared to wild-type (wt). Data for Rad21-5XFLAG ChIP and mock (non-tagged control; notag) are presented as RPM values. RPM, Reads Per Million mapped reads. **(B)** Metaplot (top) and heatmap (bottom) analysis of Rad21 ChIP signals at convergent intergenic regions (n = 292) in indicated mutants. **(C)** Box plots showing normalized Rad21 intensity (Rad21 IP normalized by mock) for the regions ± 300 bp from the Rad21 peaks as in (B). **(D)** Genome browser showing a comparison of Rad21 distribution along a 3,310 – 3,520 kb region of chromosome II. Only convergent gene pairs (forward-strand gene, blue box; reverse-strand gene, red box) overlapped with Rad21 peaks are indicated below. Box plots represent the median (center bold line), 25th and 75th percentiles (box plot boundaries), and the whiskers extending to ±1.5× interquartile range (IQR).

To further examine our results, we performed Rad21 ChIP-seq in *cid14*Δ and *exo2*Δ mutants. Cid14 is a component of the Cid14/Trf4-Air1-Mtr4 polyadenylation (TRAMP) complex, which facilitates RNA turnover by stimulating exosome activity and contributes to faithful chromosome segregation and establishment of sister chromatid cohesion during the S phase (Callahan and Butler, 2010; Win et al., 2006). In *cid14*Δ cells, we found a notable decrease in Rad21 levels (Figure 1 – figure supplement 1G-H). In contrast, Rad21 occupancy was unchanged in a mutant of Exo2 (*exo2*Δ) (Figure 1 – figure supplement 1G-H), which encodes the *S. pombe* homolog of the XRN1 5’-3’ exonuclease (Szankasi and Smith, 1996), indicating that the defect in cytoplasmic 5’-end RNA decay does not affect the Rad21 occupancy. While cohesin complex is preferentially enriched at convergent genes and evenly distributed throughout the genome in interphase, pol III-transcribed tRNA is the second most cohesin-enriched region and showed distinct cohesin peaks (Figure 1 – figure supplement 1I). Nuclear exosome and TRAMP complex are also well-known to process various RNA substrates, including tRNA, snoRNA, or rRNA. In *rrp6Δ* or *cid14Δ* mutants, but not in *pfs2-11* mutant cells, Rad21 is indeed disrupted at these tRNA loci, supporting the direct involvement of RNA processing factors in regulating cohesin occupancy at not only in Pol II-dependent genes but also in polI/III-dependent genes in a genome-wide manner.

In *S. pombe*, Swi6^HP1^ recruits Rad21 to heterochromatin regions by physically interacting with the cohesin complex (Bernard et al., 2001). Consistently, we found by Rad21 ChIP-seq that Rad21 levels at the centromeres and *tlh1^+^* subtelomeric regions (*tel1R*) decreased in *swi6*Δ cells (Figure 1 – figure supplement 2A). However, as reported previously (Bernard et al., 2001; Schmidt et al., 2009), a lack of Swi6^HP1^ had no discernible effects on the Rad21 distribution at the chromosome arms (Figure 1– figure supplement 1G-H). Thus, this indicates that Rad21 occupancy at the euchromatic chromosome arms is regulated by mechanisms distinct from those at heterochromatin regions.

We then checked whether Rrp6 and related factors were required for cohesin localization at the heterochromatin regions such as the centromere and *tlh1^+^* subtelomeric regions. Unlike the results obtained from euchromatin, in *rrp6*Δ and *pfs2-11* cells, we observed no discernible changes in Rad21 enrichment at heterochromatin (Figure 1 – figure supplement 2A) except at tRNA locus which resided within the heterochromatin. On the contrary, in *cid14*Δ and *dis3-54* cells, we found a reduced Rad21 occupancy at heterochromatin, particularly at subtelomeres (Figure 1 – figure supplement 2A), indicating the distinct function of these factors in regulating subtelomere silencing. Moreover, all of the aforementioned changes in Rad21 ChIP-seq intensities were not due to changes in the Rad21 expression (Figure 1 – figure supplement 2B). Altogether, our results emphasize the novel regulatory role of 3’- end processing, which enables the stable and proper positioning of cohesin across the chromosome arm, especially at the 3’-end of convergent genes (Figure 1D).

### Rrp6 directly regulates cohesin occupancy

Rrp6 is also known to play a central role in mediating proper termination by interacting with various cofactors (Fox and Mosley, 2016). To evaluate the direct regulation of cohesin localization by 3’-end RNA processing and transcription termination, we focused on Rrp6 to dissect the involvement of Rrp6 in regulating cohesin dynamics and turnover. We temporally depleted Rrp6 and then rescued/restored using a tetracycline-regulated system in a time-course manner (Zilio et al., 2012) (Figure 2A-B). We monitored PK-tagged Rad21 occupancy and discovered that Rad21 enrichment at converging intergenic sites (Figure 2C-E) was gradually decreased upon *rrp6^+^* knockdown (-ahTet 8 and 12 hours). When we rescued *rrp6^+^* expression by adding ahTet (+ahTet 0.5 and 4 hours), Rad21 occupancy was quickly restored (Figure 2C-E). Thus, the kinetics of *rrp6^+^* expression was highly correlated with the binding intensity of Rad21, indicating that the Rrp6-dependent regulation of Rad21 is reversible and that stabilization of the cohesin complex requires Rrp6.

**Figure 2.**
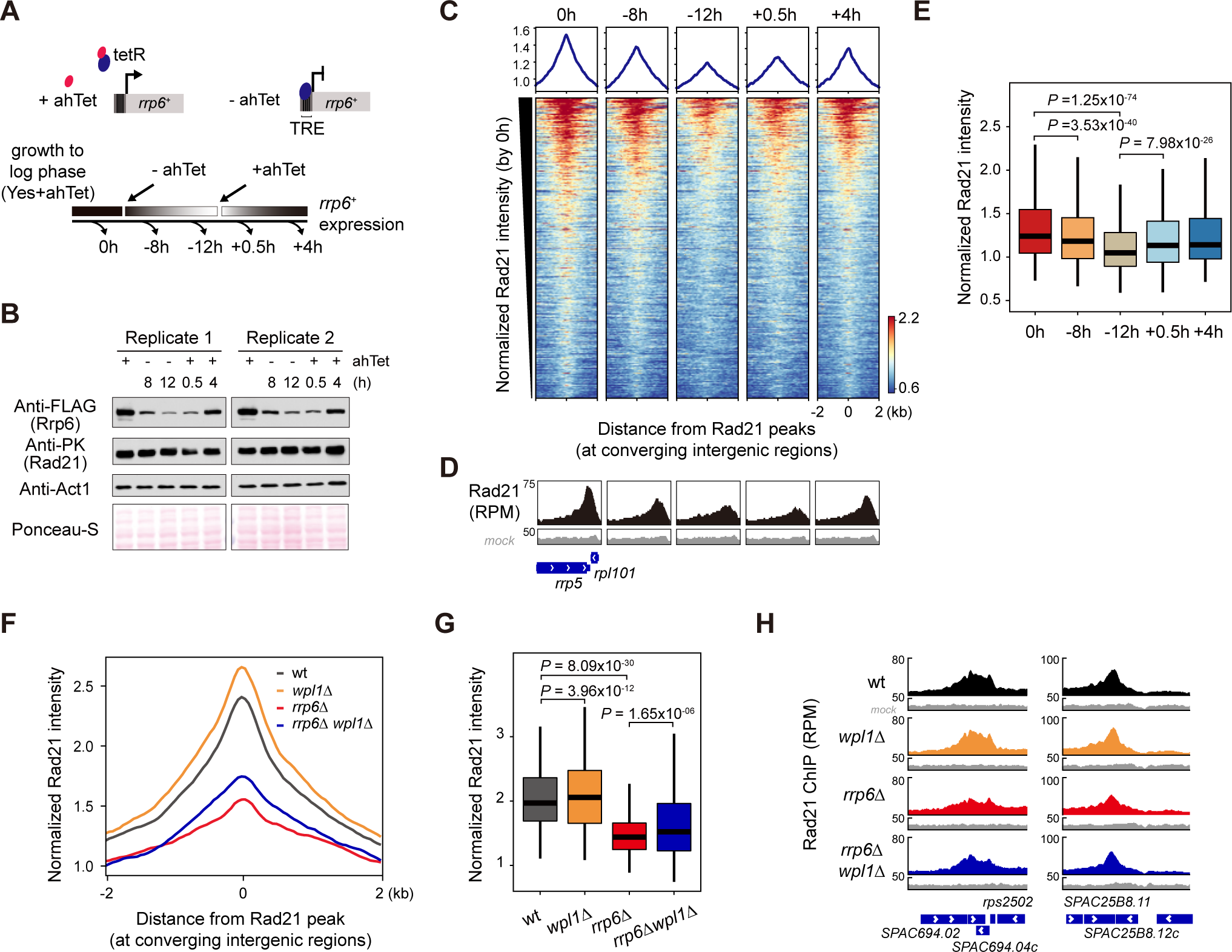
Rrp6 directly regulates cohesin occupancy. **(A)** Scheme of a tetracycline-regulated system for temporally controlled regulation of *rrp6*^+^ gene expression; ahTet indicates the tetracycline analog, anhydrotetracycline. **(B)** Western blot validating the successful knockdown and rescue of FLAG-tagged Rrp6 expression. **(C, D)** Heatmaps (C) showing the distribution of normalized Rad21 signals (Rad21 IP normalized by mock) across a time-course of ahTet withdrawal and treatment, and representative genome browser results (D) showing Rrp6-dependent changes in Rad21 occupancy. **(E)** Box plot showing normalized Rad21 intensity (Rad21 IP normalized by mock) changes across a time-course of ahTet withdrawal and treatment. **(F, G)** Metaplot (F) and genome browser results (G) showing Rrp6- and Wpl1-dependent changes of Rad21 occupancy at converging intergenic regions (n = 292). **(H)** Representative genome browser results showing Rrp6/Wpl1 dependent changes of Rad21 at convergent gene pairs.

Wapl, one of the regulatory subunits in the cohesin complex, releases cohesin from chromatin (cohesin is stabilized on the chromatin when Wapl is depleted) and restricts chromatin loop extension (Dauban et al., 2020; Haarhuis et al., 2017; Kueng et al., 2006; Vernì et al., 2000; Wutz et al., 2017). In fission yeast, we examined the interplay between Wapl and Rrp6 in cohesin turnover by combining the deletion mutant of the *wpl1^+^* with *rrp6*Δ (Figure 2 – figure supplement 1A). In *wpl1*Δ cells, we detected increased cohesin residence at converging intergenic regions, as compared to wt cells (Figure 2F-H). Notably, the reduced binding of Rad21 in *rrp6*Δ cells was partially rescued (*P value*=1.65 x 10^-06^) in the *rrp6*Δ *wpl1*Δ double mutant (Figure 2G). These results suggest that Rrp6 is directly involved in governing the turnover of cohesin at converging intergenic sites.

Along with generating contact domains in the G1 phase of the cell cycle (Bernard et al., 2008; Mizuguchi et al., 2014; Tanizawa et al., 2017), cohesin mediates sister chromatid cohesion following the S phase (Haering et al., 2008; Losada et al., 1998; Tóth et al., 1999). Since our results imply a direct linkage between Rrp6 and cohesin occupancy, we speculated that Rrp6 might play a role in the establishment of sister chromatid cohesion along chromosome arms. To prove this, we examined the state of sister chromatid cohesion using strains expressing LacI-GFP bound to LacO repeats inserted at *ade6*^+^ sites linked to the chromosome arm (Nabeshima et al., 1998; Yamamoto and Hiraoka, 2003). Wild-type and *swi6*Δ cells usually displayed a single spot of GFP, indicating normal sister chromatid cohesion along the chromosome arms (Figure 2 – figure supplement 1B-C). In contrast, *rrp6*Δ cells exhibited a higher proportion of dual GFP foci at the *ade6^+^* locus, indicating a defect in arm cohesion (Figure 2 – figure supplement 1B-C). Since Swi6 plays a major role in the centromere-specific accumulation of Rad21, we hypothesized that combining *rrp6*Δ and *swi6*Δ would show a synergistic effect. As expected, we observed a synthetic growth defect in the *rrp6*Δ *swi6*Δ double mutant, indicating that the function of Rad21 was synergistically disrupted in these cells (Figure 2 – figure supplement 1D). These data provide an additional clue that Rrp6 is involved in the regulation of sister chromatid cohesion via regulating cohesin occupancy at the chromosome arms.

### Nuclear exosome and Pfs2 regulate the higher-order 3D genome architecture via rad21

To elucidate how the loss of Rad21 occupancy upon defects in 3’-end RNA processing and transcription termination affects the 3D chromatin organization, we performed *in situ* Hi-C (Rao et al., 2014) on wt, *rrp6*Δ, *pfs2-11*, and *rad21-K1* (a loss-of-function mutant in the cohesin complex) fission yeast cells. To exclude the cell-cycle effects shown in the *rad21-K1* mutant at the restrictive temperature (37°C) (Tatebayashi et al., 1998), we conducted the Hi-C experiments in semi-permissive temperature (32°C), as we did for *pfs2-11* cells. The Hi-C experiments were performed in two biological replicates and showed high reproducibility of contact distribution according to the Pearson correlation coefficient (Figure 3 – figure supplement 1A) and of Hi-C interaction matrices using HiCRep (Figure 3 – figure supplement 1B). Consistent with previous studies in fission yeast (Mizuguchi et al., 2015, 2014), we observed centromere-proximal arm-arm interactions and Rabl chromosome configuration (Dong and Jiang, 1998; Funabiki et al., 1993; Jin et al., 2000), which is a polarized array of interphase chromosomes such as centromere clustering, in wt cells (Figure 3 – figure supplement 1C). We also confirmed, consistent with previous finding (Mizuguchi et al., 2014), that the interaction domains observed in wt cells were abolished in *rad21-K1* cells (Figure 3A, Figure 3 – figure supplement 2A). Together, these results confirm the central role of cohesin in shaping global chromosome territories in fission yeast and validate our application of Hi-C.

**Figure 3.**
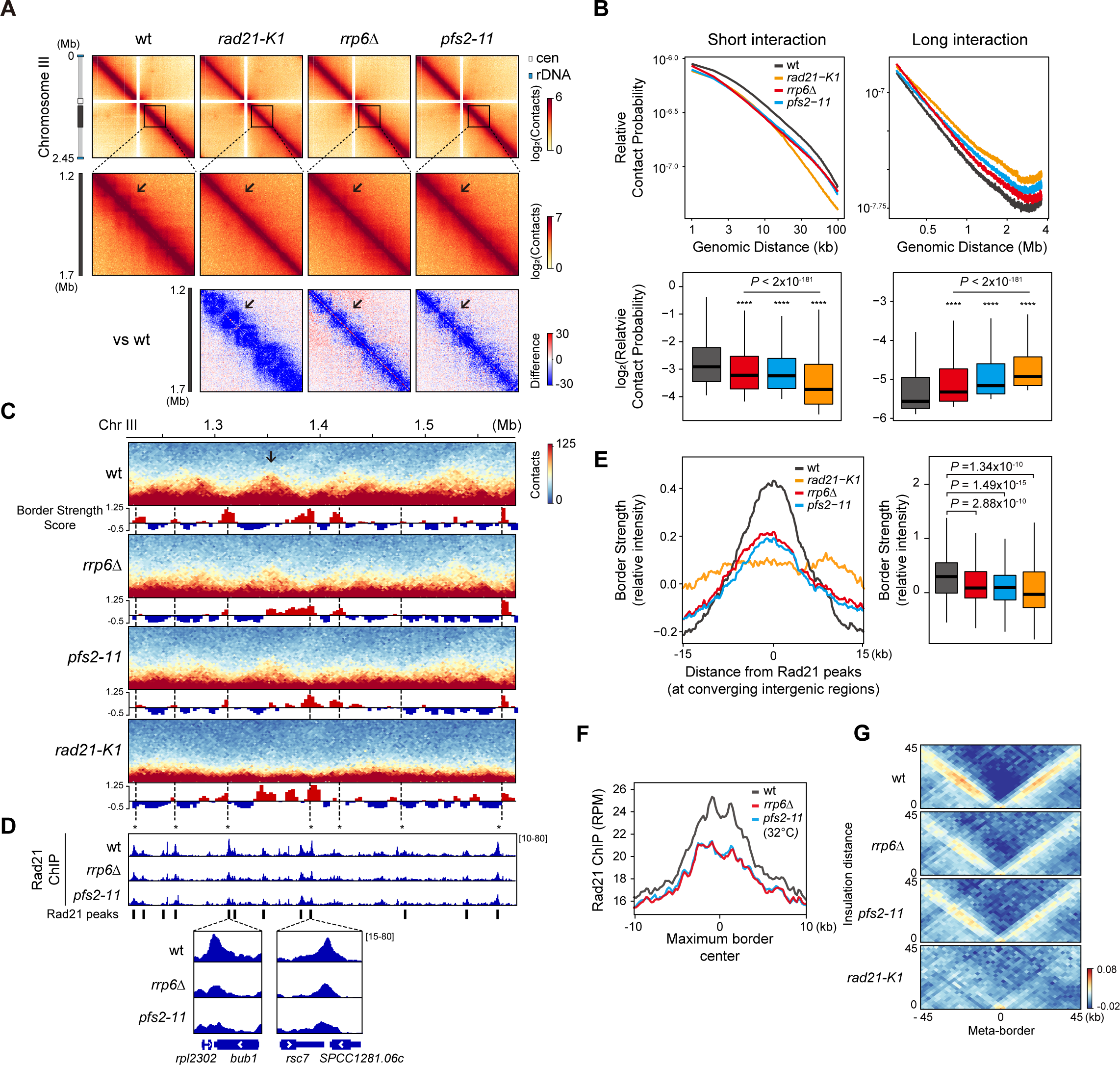
Nuclear exosome and Pfs2 regulate the higher-order 3D genome architecture via rad21. **(A)** *In situ* Hi-C map (3-kb resolution) showing the Hi-C interactions in chromosome III (top) and the 1,200 - 1,700 kb region of chromosome III (middle) of wt, *rad21-K1, rrp6*Δ, and *pfs2-11*. The relative Hi-C map (bottom) comparing mutants (*rad21-K1, rrp6*Δ, and *pfs2-11*) with wt was calculated by subtraction (mutant – wild type). Arrows indicate notable changes in the interaction domains. **(B)** Relative contact probability (RCP) plot (top) showing contact frequency relative to genomic distance. Box plots (bottom) comparing the contact probabilities of short-range (< 100-kb) and long-range (> 300-kb) interactions in indicated mutants. *P* values were calculated using the paired Wilcoxon rank- sum test. **(C)** Representative Hi-C map showing the 1,200 - 1,700 kb region of chromosome III (3-kb resolution). Border Strength is calculated with 3-kb resolution of 15-kb window sizes. See also Figure 3 – figure supplement 2C. **(D)** Rad21 ChIP-seq (RPM) of wt, *rrp6*Δ, and *pfs2-11* track is aligned with the above Hi-C map (C). In particular, sites of reduced border strength along with reduced Rad21 peaks upon defects in Rrp6 or Pfs2 are marked with asterisks. The lower panels illustrate a zoomed-in view of Rad21 ChIP. **(E)** Metaplot (Left) and boxplot (Right) of border strength at Rad21 peaks at converging intergenic sites. **(F)** Enrichment of Rad21 ChIP at the center of domain border in wt, *rrp6*Δ, and *pfs2- 11* mutants. **(G)** Aggregate plots generated from Hi-C data for Rad21 peaks (n = 292) showing discrete cohesin-mediated border. Box plots represent the median (center bold line), 25th and 75th percentiles (box plot boundaries), and the whiskers extending to ±1.5× interquartile range (IQR).

In our Hi-C analysis of *rrp6*Δ and *pfs2-11* cells, we observed that similar to the *rad21-K1* cells, there was a significant decrease in intra-chromosomal (*cis*) arm interactions (Figure 3A, Figure 3 – figure supplement 2A). Next, we compared the contact probability decay according to the genomic distance, reflecting the general chromatin folding state. The *rrp6*Δ and *pfs2-11* mutant cells exhibited a quantitative reduction of short-range interactions within 100-kb (kilobase), with an increase in long-range (>300-kb) interactions (Figure 3B, Figure 3 – figure supplement 2B). Both changes in short- and long-range interactions were reminiscent of the patterns observed in *rad21-K1* cells (Figure 3B, Figure 3 – figure supplement 2B). Together, these results demonstrate that 3’-end RNA processing factors, including Rrp6 and Pfs2, are required to maintain 3D interaction domains and local compaction across the intra-chromosomal arms by modulating cohesin occupancy.

To further dissect how RNA processing factor–associated loss of cohesin binding affects the border formation, we calculated the border strength score, which represents a potential border as previously described (Kim et al., 2016; Van Bortle et al., 2014) whereby the border strength is quantified by a relative ratio of two adjacent intra-domain with inter-domain contact frequencies at a resolution of 3-kb with a window size of 15-kb (Figure 3 – figure supplement 2C). As expected, in wt cells, we detected a high border strength separating the 3D interaction domains (Figure 3C), and it coincided with Rad21 ChIP-seq peaks enriched at converging intergenic sites (Figure 3C-D). Using the border strength, we estimated the size of wt interaction domains in the median size of 65-kb, consistent with the previous findings (Mizuguchi et al., 2014; Tanizawa et al., 2017). For further validation, we compared the border strength scores derived from two wt Hi-C datasets (ours and GSM2446268) and observed that both were similarly high at the Rad21 peaks (Figure 3 – figure supplement 2D). We next plotted the border strength scores according to the Rad21 levels in wt cells and found a positive correlation between the border strength and Rad21 occupancy (Figure 3 – figure supplement 2E). We also analyzed the published Psc3 (another cohesin subunit) ChIP-chip (GSM1370192) and found that Psc3 was enriched at the maximum border center, defined by the region with maximum border strength scores across the genome (Figure 3 – figure supplement 2F). Thus, these results validate our border strength scores and indicate that strong domain borders at converging intergenic sites are occupied by enriched cohesin.

Using this validated approach for quantifying border strength, we examined domain border strength in *rrp6*Δ and *pfs2-11* cells and detected a significant reduction, similar to what is observed in *rad21-K1* cells (Figure 3C,E). This reduction coincided with a decrease in cohesin occupancy (Figure 3D). The degree of border strength changes in each mutant (Figure 3E) indeed corresponded with the changes in Rad21 occupancy at the borders (Figure 3F). To confirm this in a genome-wide fashion, we visualized the Hi-C interactions near Rad21 peaks using insulation distance as previously described (Vara et al., 2019). As expected, we observed that inter-domains (located between adjacent intra-domains, Figure 3 – figure supplement 2C) exhibited low Hi-C contacts in wt cells (Figure 3G), suggesting that enriched Rad21 sites insulate adjacent intra-domains. Notably, similar to *rad21-K1* cells, we observed increased Hi-C contacts within inter-domains in *rrp6Δ* and *pfs2-11* cells, compared to those in wt cells (Figure 2G).

To further examine whether the nuclear exosome indeed affects the 3D chromatin structures, we additionally performed *in situ* Hi-C using defective mutants in Dis3 (*dis3-54*), another core and catalytic subunit of the nuclear exosome, with high reproducibility (Figure 3 – figure supplement 3A-B). A cold-sensitive mutant of Dis3, *dis3-54*, shows a reduced RNase activity at its restrictive temperature (20°C) (Mukarami et al., 2007). As with Pfs2 and Rad21, Dis3 is essential for viability, and thereby we grew *dis3-54* cells at permissive (wt at 34°C) and semi-permissive temperature (defect at 30°C). Consistent with *rrp6*Δ cells, we observed a decrease in intra-chromosomal (*cis*) arm interactions (Figure 3 – figure supplement 3C) and short-range interactions (Figure 3 – figure supplement 3D) in *dis3-54* defect cells, yet albeit weakly due to the use of partially loss-of-function mutant. We compared the changes in the border strength (Figure 3 – figure supplement 3E-F) and the insulation distance analysis (Figure 3 – figure supplement 3G) in *dis3-54* wt and mutant cells, suggesting a contribution of Dis3 in the 3D chromatin organizations. Together, these findings support a regulatory role for nuclear exosome in shaping the proper domain borders through the regulation of cohesin positioning.

As Rad21-enriched converging intergenic sites showed discrete domain boundaries, we next wondered whether other Rad21 strongly-enriched loci, including tRNA, rRNA, or promoters, also form cohesin-mediated domain boundaries. However, the intensity of the border strength at each group was relatively smaller than the converging intergenic sites. In particular, tRNA loci distributed across the chromosome arms were strongly enriched with cohesin, and the cohesin localization was dependent on nuclear exosome and Cid14 (TRAMP complex) (Figure 1 – figure supplement 1I). Nevertheless, the border strength at the tRNA loci was not as strong as the border strength at converging intergenic sites and was not affected by the deletion of Rrp6 or by *rad21-k1* mutant cells (Figure 3 – figure supplement 2G). Therefore, our findings indicate that proper positioning of cohesin at 3’-end of convergent genes via RNA processing factors ensures the formation of discrete and strong domain boundaries.

### Proper transcription termination coupled with 3’-end RNA processing is important for the establishment of cohesin at the 3’-end of convergent genes

Previously, it was suggested that Rad21 is pushed towards the 3’-end of convergent genes by the elongating transcription machinery (Busslinger et al., 2017; Gullerova and Proudfoot, 2008; Heinz et al., 2018; Lengronne et al., 2004). To investigate this in a genome-wide manner, we analyzed transcription levels, distribution of Rad21 and RNA Pol II Ser2 CTD phosphorylation (Ser2P), and border strength using mRNA-seq, ChIP-seq, and *in situ* Hi-C. In wt cells, we first sorted 1270 pairs of convergent coding genes by transcription strength and then plotted the distribution of Ser2P and Rad21 along with border strength in the same order (Figure 4A, Figure 4 – figure supplement 1A). We observed a strong enrichment of Ser2P and Rad21, coinciding with high border strength (Figure 4A), suggesting that Rad21 accumulation at highly transcribed gene pairs is responsible for forming domain borders in fission yeast.

**Figure 4.**
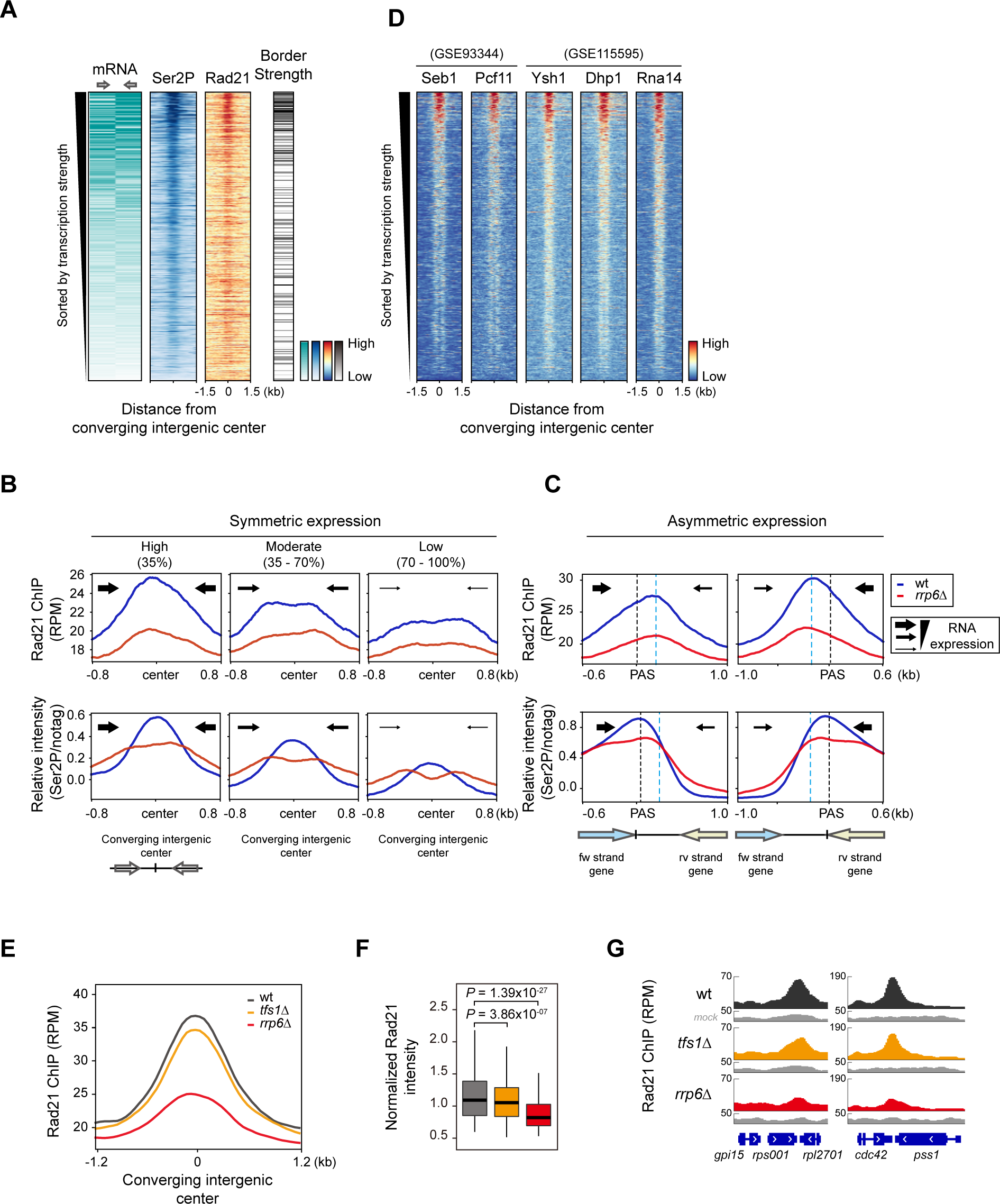
Proper transcription termination coupled with 3’-end processing is important for the establishment of cohesin at the 3’-end of convergent genes. (**A**) Heatmaps sorted by the median mRNA expression (RPKM) of total convergent gene pairs. ChIP- seq of Rad21 (RPM) and RNAPII Ser2P phosphorylation (Ser2P) and the border strength were plotted against the total converging intergenic center. (**B, C**) Metaplot analysis of Rad21 (RPM) and Ser2P (normalized by FLAG-immunoprecipitated non-tagged control; notag) occupancy at either symmetrically (B) or asymmetrically (C) expressed pairs of convergent genes. Converging gene pairs with symmetric (mRNA RPKM difference between the gene pairs is less than 2-fold) or asymmetric transcription strength are sorted by mRNA RPKM and divided into either three groups (highly expressed (n = 148); 0 - 35%, moderately expressed (n = 152); 35 - 70%, and lowly expressed (n = 130); 70 -100%) (B) or two groups (forward convergent genes strongly transcribed than reverse convergent genes (n = 190) (Left) and vice versa (Right) (n = 249)) (C). Blue dotted lines indicate the average Rad21 enrichment center. **(D)** Heatmaps sorted by the same order as in (A) showing the co-localization of Rad21 and Ser2P with termination factors (Dhp1, Seb1) and CPSF complex (Pcf11, Rna14, and Ysh1) at highly transcribed converging genes. Transcription termination factors (Pcf11, Seb1, Rna14, Ysh1, Dhp1) were derived from published ChIP-seq data (GSE93344 and GSE 115595). **(E, F)** Metaplot (E) and box plot (F) of Rad21 peaks (normalized by mock) in indicated mutant cells. **(G)** Representative genome browser results showing Rad21 enrichment in indicated mutants (*tfs1*Δ and *rrp6*Δ) at convergent gene pairs.

We considered that RNAPII accumulated in the intergenic region of the convergent genes could be a trace of RNAPII backtracking activity and a recent study also reported that backtracking of RNA polymerase induces the accumulation of condensin (Rivosecchi et al., 2021). Consistent with a previous study in mammalian cells (Busslinger et al., 2017), at symmetrically expressed sites, Rad21 was positioned symmetrically, and this correlated with the accumulated Ser2P signals at the same converging intergenic sites. Given that the regions we were interested in were highly transcribed loci and may be interfered with the background immunoprecipitation, we normalized Ser2P signals with mock (notag) immunoprecipitation (De Almeida et al., 2010; Pokholok et al., 2005). Deletion of Rrp6 reduced the elongating RNAPII signals at the 3’-ends and increased transcriptional readthrough (Figure 4B, Figure 4 – figure supplement 1A, see red boxes), which indicated the defects in transcription termination. At asymmetrically expressed pairs of convergent genes, we also found the reduced RNAPII at polyadenylation sites (PAS) and downstream-shifted Ser2P signals (Figure 4C). This indicates that reduced cohesin binding at the 3’-end of convergent genes may be linked to Rrp6-mediated termination defects. Recent reports about the novel role of nuclear exosomes in promoting transcription termination by targeting the 3’-end of RNAs exposed from backtracked RNAPII also supported our findings.

Transcription termination is directly linked to mRNA processing machinery, which occurs co-transcriptionally. We further investigated the enrichment of 3’-end processing machinery (Pcf11, Ysh1/CPSF73, and Rna14/CSTF3) and transcription termination factors (Seb1 and Dhp1/Xrn2) using published ChIP-seq data (Larochelle et al., 2018; Wittmann et al., 2017). We also found that these factors were also co-enriched with Ser2P and cohesin at converging intergenic centers (Figure 4D). We observed that Rad21 levels were decreased proportionally to the level of transcription termination factors in *rrp6*Δ cells, compared to wt cells (Figure 4 – figure supplement 1B-C). Together, these data indicate that cohesin and transcription termination-related factors are enriched at highly transcribed convergent gene pairs, correlated with the strong domain borders.

The transcription factor TFIIS, stimulates the intrinsic cleavage activity of RNAPII, thereby allowing the backtracked RNAPII to resume elongation (Izban and Luse, 1992; Reines, 1992). In TFIIS- deleted cells, premature dissociation of RNAPII occurs, resulting in premature termination (Lemay et al., 2014; Sheridan et al., 2019; Sigurdsson et al., 2010). Compared to *rrp6Δ*, when we deleted Tfs1, we only observed a very slight decrease of Rad21 enrichment at converging intergenic sites. It is reported that although when TFIIS is absent, an intrinsic cleavage activity of RNAPII slowly allows the reactivation of RNAPII (Sigurdsson et al., 2010). As cohesin is translocated by elongating Pol II from promoters and once it reaches toward the 3’-ends, the interplay between RNAPII backtracking and transcription termination (linked with 3’-end processing machinery and nuclear exosomes) may play a crucial role in stabilizing the cohesin complex.

We further explored how the readthrough transcripts produced by the loss of Rrp6 (Figure 4 – figure supplement 1D) affect the localization of Rad21. To see whether there is a negative correlation between readthrough transcripts and cohesin intensity (increased readthrough transcripts may cause a greater decrease in cohesin level), we calculated the termination index (Figure 4 – figure supplement 1E). In *rrp6Δ* cells, we found that changes in Rad21 level in converging intergenic regions showed no negative correlations with the changes in the level of transcription readthrough produced from the pair of convergent genes (Figure 4 – figure supplement 1F). We conclude that the readthrough transcripts produced from defective RNA processing factors do not solely detach cohesin from the binding sites.

### Cohesin is redistributed to facultative heterochromatin at retrotransposon upon Rrp6 depletion

We revisited the Rad21 changes upon deletion of Rrp6 to see whether Rad21 is redistributed across the genome. We merged 701 Rad21 peaks from 875 and 719 Rad21 peaks in wt and *rrp6Δ* cells, respectively. Interestingly, we observed 123 increased/newly-created Rad21 peaks (17 %) in *rrp6Δ* cells, which were nearly non-detectable in wt cells (Figure 5A, a bold red line). Furthermore, 51% of these new Rad21 peaks were localized in gene bodies (Figure 5 – figure supplement 1A), where 80% of these genes were either Tf2 retrotransposon elements or proximal to these elements. In *S. pombe*, there is only one family of full-length long terminal repeat (LTR) retrotransposon element (Tf2) (Bowen et al., 2003) with an average size of 4-kb. At all 13 Tf2 elements, we detected substantially increased Rad21 levels in *rrp6Δ* cells (Figure 5B, left), while Rad21 levels did not change in pfs2-11 cells, compared to wt cells (Figure 5B, right). Additionally, we combined Wpl1 deletion (*wpl1Δ*) with *rrp6Δ* and observed more dramatically increased Rad21 levels in *wpl1Δ rrp6Δ* cells than those in wt, *rrp6Δ*, and *wpl1Δ* cells (Figure 5 – figure supplement 1B), confirming that Rrp6/Wpl1 independently regulates cohesin positioning. Thus, these data indicate that upon Rrp6 deletion, cohesin is redistributed to specific genomic loci (Tf2).

**Figure 5.**
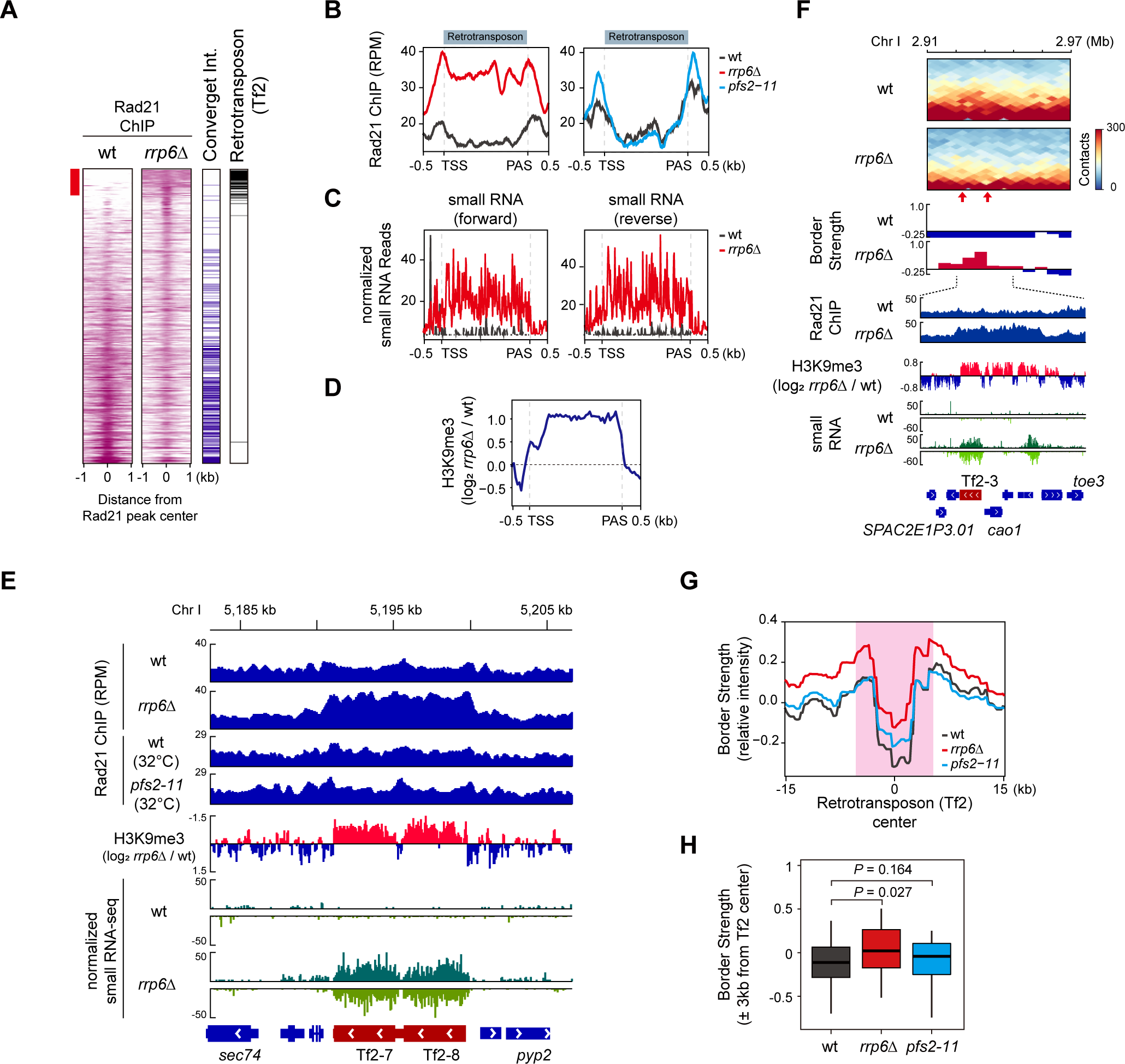
Cohesin is redistributed to facultative heterochromatin at retrotransposon upon Rrp6 depletion. **(A)** Heatmaps of Rad21 ChIP-seq signals (RPM) at combined Rad21 peaks at both wt and *rrp6*Δ, sorted by change (*rrp6*Δ/wt) in Rad21 intensity. Assignment of Rad21 peaks that overlap or proximal to either convergent intergenic regions (Convergent Int) or Retrotransposon (Tf2 elements) are shown on the right panel. **(B)** Metaplots of Rad21 (RPM) binding of wt, *rrp6*Δ, and *pfs2-11* at 13 retrotransposon elements. **(C, D)** Normalized (forward or reverse) small RNA reads of wt and *rrp6*Δ (C) and H3K9me3 (log2 *rrp6*Δ/wt) (D) at 13 retrotransposon elements. **(E)** Genome browser example of chromatin features at retrotransposon element Tf2-7-Tf2-8 where Rad21 binding is increased upon Rrp6 deletion. **(F)** *In situ* Hi-C map (5-kb resolution) (top) and the border strength (middle) and chromatin features (bottom) at retrotransposon element Tf2-3. **(G)** Average profile of border strength plotted against the center of 13 retrotransposon elements. **(H)** Box plot showing significantly increased border strength at the center of 13 retrotransposon elements in *rrp6*Δ compared to wt and *pfs2-11*.

Recent studies reported RNAi-dependent heterochromatin domain (HOOD) formation at repetitive DNA elements such as centromeric repeats and retrotransposons, and its formation is closely associated with Rrp6. In the absence of Rrp6, which is also known for processing/silencing heterochromatic transcripts, accumulation of siRNA and heterochromatin modifications (H3K9 methylation) were observed. To elucidate whether Rad21 redistribution at Tf2 upon Rrp6 deletion follows the same process as Rad21 recruitment at heterochromatin (Bernard et al., 2001; Nonaka et al., 2002; Reyes-Turcu et al., 2010), we performed H3K9me3 ChIP-seq and small RNA-seq in wt and *rrp6Δ* cells. Indeed, we observed elevated levels of small RNA and H3K9me3 at all Tf2 elements in *rrp6Δ* cells compared to wt cells (Figure 5C-D), consistent with previous findings (Yamanaka et al., 2013). Notably, these elevated small RNA and H3K9me3 levels coincided with Rad21 recruitment at specific Tf2 loci in *rrp6Δ* cells (Figure 5E). Among 32 defined HOOD loci, all 13 Tf2 are included within HOOD regions, yet Rad21 distribution and heterochromatin markers (small RNA and H3K9me3) on Tf2-excluded remaining HOOD regions were relatively unaffected in *rrp6Δ* (data not shown). Furthermore, published H3K9me2 ChIP-seq (GSE114540) (Parsa et al., 2018) showed no notable increase in *pfs2-11* mutant compared to wild-type (Figure 5 – figure supplement 1C). This implies that the production of small RNAs upon Rrp6 deletion induces RNAi-mediated facultative heterochromatin formation on retrotransposon and therefore redistributes Rad21 to these loci in an indirect manner.

We then asked whether Rad21-enriched Tf2 elements could function as a domain border in *rrp6Δ* cells. To this end, we calculated the border strength at Tf2 elements in wt, *pfs2-11*, and *rrp6Δ* cells. Importantly, we found increased border strength at Tf2 loci, enriched with Rad21, H3K9me3, and small RNA, in *rrp6Δ* cells compared to wt cells (Figure 5F). In contrast, loss of Pfs2 (*pfs2-11*) did not affect the formation of domain border (Figure 5G-H), probably due to unchanged Rad21 levels (Figure 5B, E) at Tf2 elements. Therefore, we concluded that Rrp6 is essential for establishing the domain borders and ensuring genome stability by regulating proper cohesin positioning.

### The role of Rrp6^Exosc10^ in regulating cohesin and 3D chromatin structures observed in *S. pombe* is similarly conserved in mESC

The functions of the exosome and the cohesin complex are highly conserved from yeast to human. To examine whether the role of the nuclear exosome in 3D genome organization is conserved in the mammalian genome, we performed *in situ* Hi-C using mouse embryonic stem cells (mESCs) with RNAi-mediated knockdown of relevant exosome subunit (Exosc10^Rrp6^) and cohesin subunit (Rad21) (Figure 6 – figure supplement 1A). Hi-C analysis achieved high reproducibility between two biological replicates (Figure 6 – figure supplement 1B). Interestingly, Rad21 knockdown (*Rad21KD*) perturbed the global 3D genome organization in a manner that was consistent with our results obtained in *S. pombe*, but knockdown of Exosc10 (*Exosc10KD*) only marginally affected global chromatin organization compared to that in *Egfp* knockdown (Control) cells (Figure 6A, Figure 6 – figure supplement 1C-E). Specifically, we observed that markers of global chromatin organization, including the relative contact probability (Figure 6A), the frequency of *cis*-interactions (Figure 6 – figure supplement 1C), and the number of contact domains/chromatin loops (Supplemental table 3), were disrupted in *Rad21KD* cells, but not in *Exosc10KD* cells, compared to Control cells.

**Figure 6.**
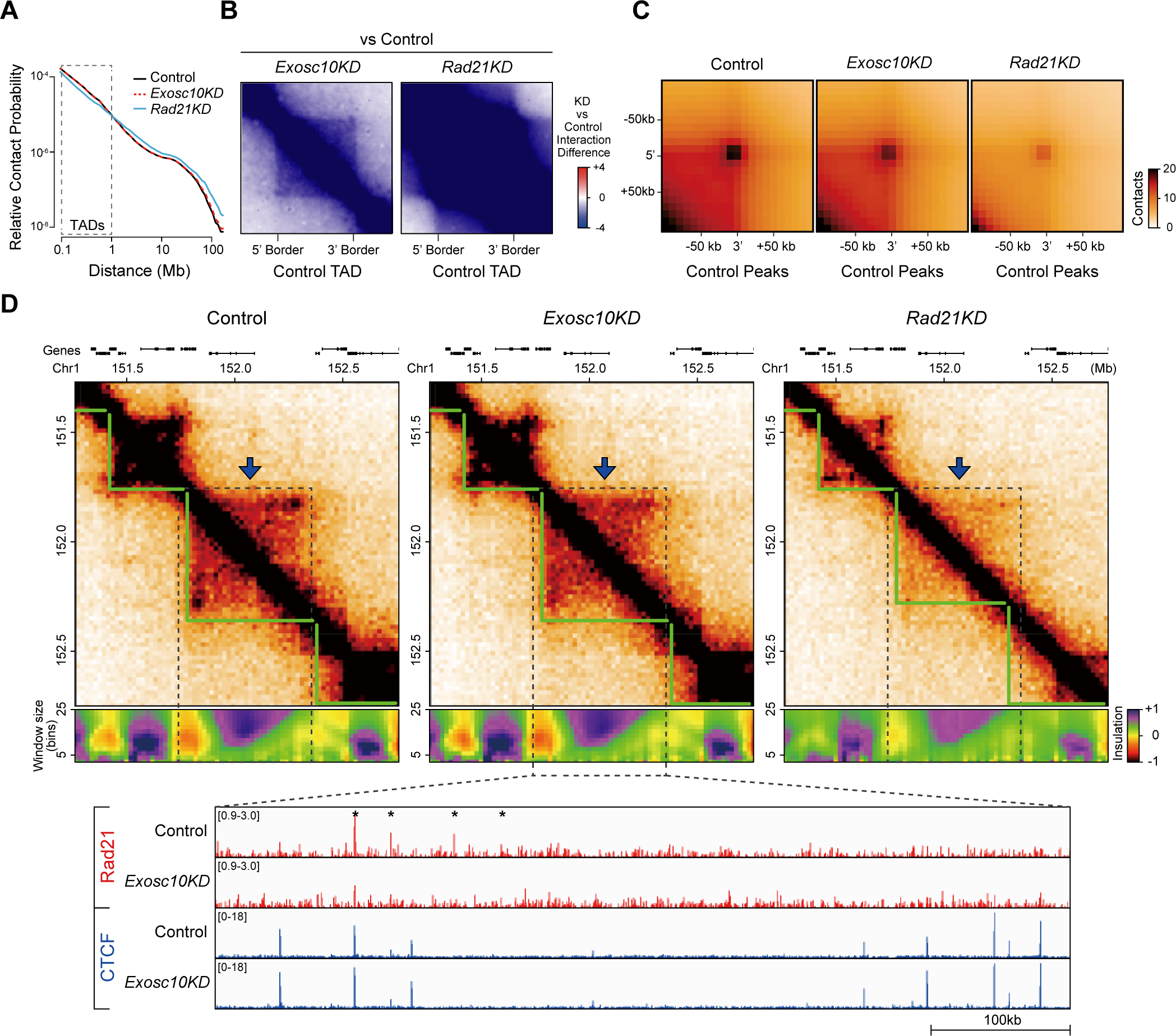
The role of Rrp6^Exosc10^ in regulating cohesin and 3D chromatin structures observed in *S. pombe* is similarly conserved in mESC. **(A)** Relative contact probability (RCP) plot showing contact frequency relative to genomic distance. The dashed box displays the size range of TADs (≤1 Mb). **(B)** Aggregate TAD analysis (ATA) comparing interactions of *Exosc10KD* and *Rad21KD* with Control (*EgfpKD*) cells. Control TADs were used for ATA (n = 4508). **(C)** Aggregate peak analysis (APA) comparing Control, *Exosc10KD*, and *Rad21KD* cells. Control peaks (chromatin loops) were used for APA (n = 12,477). **(D)** Example of a TAD that lost its interactions (dashed box with blue arrow) upon *Exosc10* and *Rad21* knockdown. Hi- C contact maps (top), insulation heatmaps (middle), and Rad21 and CTCF ChIP-seq peaks (bottom; red and blue bars, respectively) are shown for Control, *Exosc10KD*, and *Rad21KD* mESCs. The ChIP-seq tracks display zoom-in browser view of dashed box showing where a TAD loses its interactions. The decreased Rad21 ChIP-seq peaks upon *Exosc10KD* are marked with asterisks.

To further investigate the roles of Exosc10 and Rad21 in local chromatin organizations, we performed ATA/APA (Aggregate TAD/Peak Analysis) at the Control TADs/loops. Importantly, the ATA/APA results showed a gradual decrease in interactions within TADs (Figure 6B) and chromatin loops (Figure 6C, Figure 6 – figure supplement 1F) from Control in *Exosc10KD* and *Rad21KD* cells, consistent with our results obtained in *S. pombe* (Figure 3). Since CTCF (which is absent in yeast) is known to mediate transcriptional insulation, co-localize with cohesin, and contribute to Rad21 positioning (Parelho et al., 2008; Wendt et al., 2008), we performed ChIP-seq of Rad21 and CTCF in *Exosc10KD* mESCs to determine the basis of the gradual decrease in local interactions. In these cells, we observed a slight decrease in Rad21 occupancy at CTCF peaks near TAD boundaries (*P* value= 0.006), whereas the CTCF peaks showed marginal changes (Figure 6D, Figure 6 – figure supplement 1G). Interestingly, we observed reduced Rad21 occupancy irrespective of CTCF presence (Figure 6D, Figure 6 – figure supplement 6G), implying that even in mammalian genomes, Exosc10 can partially regulate the cohesin occupancy. Finally, we found that the local interactions within the Control TAD were gradually decreased in both *Exosc10KD* and *Rad21KD* cells, as confirmed by the ATA/APA results. Furthermore, we observed that the reduction of local TAD interactions coincided with the decreased Rad21 enrichment in *Exosc10KD* cells. These results indicate that Exosc10 plays an important role in organizing the 3D genome architecture by regulating the Rad21 localization, and this role is highly conserved from pre-metazoans to metazoans.

## Discussion

In this study, we used various proteins related to RNA surveillance to examine their role in regulating cohesin complex and 3D chromatin organization. Using *in-situ* Hi-C, we observed that ablation of RNA processing factors, including nuclear exosome and Pfs2 (CPF complex), resulted in the loss of cohesin binding and disturbed cohesin-mediated domain boundaries. These results highlight the importance of RNA processing coupled with termination in ensuring proper cohesin positioning at the domain borders where convergent genes mainly exist, thereby maintaining 3D chromatin organizations. Strictly regulated 3’-end RNA processing coupled with proper termination and 3D genome architecture are essential factors for gene expression. This study proved that two processes, which were thought to be independent of each other during transcription, were closely intertwined.

In fission yeast, the cohesin complex is preferentially enriched (∼ nearly 47%) at the 3’-end of convergent genes. 3D domain borders mainly reside at the 3’end of highly transcribed convergent genes where cohesin and transcription termination factors were co-enriched. Although the intensity of cohesin binding at other cohesin-enriched loci such as tRNA was reduced in RNA processing defective mutants, those regions did not harbor discrete and strong domain borders. Moreover, the border strength at those regions was not dependent on RNA processing factors. These findings once again highlighted the importance of well-positioned cohesin complex at converging interenic sites. In particular, defects in Rrp6 caused Rad21 redistribution to retrotransposon in an indirect manner, resulting in improper domain borders.

At converging intergenic sites, elongating polymerase from both sides continuously approaches toward the 3’-ends, and we observed the accumulation of elongating Pol II (Ser2P) at the end (Figure 4), which may be resulted from the backtracking. Defect in Pfs2, which causes extensive transcription readthrough, prevents backtracked RNAPII. Nuclear exosome targets the 3’-ends of unprotected RNAs that are produced by backtracked RNAPII and promotes transcription termination. Defects in nuclear exosome or extensive transcription readthrough prevent/escape backtracked RNAPII. Recently, a study in fission yeast reported that the RNA polymerase backtracking at gene termini might contribute to the accumulation of condensin, another SMC complex (Rivosecchi et al., 2021). Nevertheless, reduced cohesin binding upon defective 3’-end RNA processing (e.g., *rrp6Δ* or *pfs2-11*) are not solely caused by passive translocation of RNAPII, as we can see in *tfs1Δ*. Taken together, these may explain the active role of RNA processing factors in concert with proper transcription termination in stabilizing the cohesin complex at the 3’end of genes.

We assumed that the production of readthrough transcripts in *rrp6Δ* might explain the decrease of cohesin binding at chromatin. However, we could not find a direct relationship between readthrough transcript and chromatin localization of cohesin in which higher levels of readthrough transcripts produced in *rrp6Δ* do not always result in cohesin reduction at converging intergenic sites. However, it is still possible that even small amounts of readthrough transcripts may disturb cohesin positioning. For instance, Xist RNA, which regulates the silencing of one of two X chromosomes via acting as a scaffold protein, binds with cohesin and causes repulsion of the cohesin from the inactivated X chromosome (Minajigi et al., 2015). SA1 and SA2, subunits of the cohesin complex, were reported to bind with various kinds of RNA substrates such as ssRNA, dsRNA and RNA:DNA hybrids (Pan et al., 2021). These evidence leaves a possibility that readthrough transcripts may play a role in regulating cohesin localization.

Furthermore, our results may provide a clue to solving the mystery of chromosomal segregation defects. Previous studies have found chromosomal segregation defects in cells mutant for transcription termination and RNA processing factors, but the underlying mechanisms connecting these have not been resolved (Graham et al., 2009; Mukarami et al., 2007; Wang et al., 2005). A recent study showed that in budding yeast, convergent genes exist at pericentromeres and act as a border that captures cohesin (Paldi et al., 2020). These findings suggest that proper cohesin localization at the pericentromeres may be necessary for chromosome segregation. Our results showed that Rrp6 deletion causes three events: 1) reduction in cohesin occupancy at the 3’-end of convergent genes, 2) increased transcriptional readthrough (defects in transcription termination), and 3) weakness in chromosome arm cohesion. Regarding these, our data suggest that Rrp6-mediated transcription termination and cohesin positioning may play an crucial role in genome integrity (chromosomal segregation).

Influenza A virus (IAV) infection induces global RNAPII transcription termination defects (Zhao et al., 2018). Influenza NS1 protein, in particular, targets the 3’-end processing machinery (CPSF30) and inhibits PAS-dependent termination. As such, IAV infection causes readthrough transcription, disrupts cohesin binding, and therefore results in chromatin decompaction. Therefore, in agreement with recent findings, our results extend the role of RNA surveillance to control the dynamics of cohesin biology and chromatin architecture. We hope that our findings will contribute to the elucidation of cohesin-mediated regulation of 3D chromatin structures, with potential implications for understanding cohesin disruptions in human cancers and cohesinophaties.

## Acknowledgements

We would like to thank M. Yanagida, K. Ekwall, C.J. Norbury and R. Allshire for providing fission yeast strains. This work was supported by a National Research Foundation of Korea Grant funded by the Ministry of Science and ICT (MSIT) (2018R1A5A1024261, SRC), and the Collaborative Genome Program for Fostering New Post-Genome Industry of the National Research Foundation (NRF) funded by the MSIT (2018M3C9A6065070).

## Author Contributions

Y. Choi, Y. Chun and D. Lee devised the concepts and designed the experiments. S. Han, T. Kim and Y. Chun analyzed the NGS data. Y. Choi and Y. Chun performed most of the experiments, including Hi-C and ChIP-seq. S. Han, and Y. Chun wrote the manuscript under the mentorship and technical supervision of D. Lee and Y. Choi.

## Supplemental Information

Supplemental Information is available for this paper.

## Materials and Methods

### Yeast strains and culture

The strains used in this study are listed in Supplemental Table 1. All deletion strains and tagged strains were constructed by gene targeting using PCR-based methods (Bähler et al., 1998). Yeast cells were grown in YES (Yeast-Extract with Supplements) medium at 30°C. The temperature-sensitive mutants, including *rad21-K1* (32°C), *pfs2-11* (32°C), and *dis3-54* (30°C) were cultured at their semi-permissive temperatures. For the *in situ* Hi-C experiments, yeast cells were grown at 32°C unless otherwise indicated.

For *dis3^+^* and *rrp6*^+^ gene knockdown and rescue experiments, their gene promoters were replaced with a tetracycline-responsive promoter harboring TRE (tetracycline response element) (Zilio et al., 2012).

Cells were first grown to mid-log phase (∼OD600 0.6) at 30°C in YES medium supplied with 2.5μg/ml of anhydrotetracycline (ahTet). To deplete Dis3 and Rrp6, cells were shifted to YES medium lacking ahTet and incubated for indicated times (∼10hr for Dis3 and 8, 12 hr for Rrp6). To recover Rrp6 expression, the cells were then shifted to fresh YES medium containing ahTet (2.5 μ g/ml) and incubated for the indicated times (0.5 and 4 hours).

### Mouse embryonic stem cell culture

E14Tg2a mESCs were cultured on feeder-free gelatin-coated plates in Glasgow minimum essential medium (GMEM, Gibco) supplemented with 10% knockout serum replacement (Gibco), 1% defined fetal bovine serum (FBS, Hyclone), 1% non-essential amino acids (NEAA, Gibco), 1% sodium pyruvate (Gibco), 0.5% antibiotic-antimycotic (penicillin-streptomycin containing, Hyclone), 0.1 mM β-mercaptoethanol (Gibco), and 1000 U/ml leukemia inhibitory factor (LIF, Millipore). E14Tg2a mESCs were grown at 37°C in air containing 5% CO2.

### RNA interference

Silencer® Selects s78573 (5 nmol, Thermo) and Silencer® Select s72659 (5 nmol, Thermo) were used as the siRNAs against Exosc10^Rrp6^ and Rad21, respectively, while the control siRNA sequence against EGFP, GUUCAGCGUGUCCGGCGAGUU, was produced by Bioneer (Daejeon, Korea). E14Tg2a mESCs (1x10^5^ cells/well) were seeded to 6-well plates. After 24 hours, the cells were transfected with 50 nM of siRNA against Exosc10, Rad21, or Egfp using DharmaFECT 1 (T-2001-03, Dharmacon) and Opti-MEM (Gibco) according to the manufacturer’s protocols. At 48 hours post-transfection, the cells were harvested and analyzed by RT-qPCR.

### In situ Hi-C

The *in situ* Hi-C library construction in fission yeast was performed as previously described (Rao et al., 2014), with slight modifications. Briefly, 2.4x10^9^ mid-log phase cells were fixed with 3% formaldehyde for 15 min and then quenched with 125 mM glycine. The fixed cells were washed with ice-cold TBS (20 mM Tris-HCl pH 7.5, 150 mM NaCl), nuclei were extracted by zymolyase digestion, and 6 μg of pelleted nuclei were used for Hi-C library construction. Each nuclear pellet was incubated in 50 μl of 0.5% SDS at 62°C for 10 minutes and then immediately quenched with 170 μl of 1.47% Triton X-100 at 37°C for 15 minutes. Chromatin was digested with 100 units of MboI (NEB) at 37°C for 2 hours and subsequently incubated at 62°C for 20 minutes to inactivate the MboI. To fill in the overhangs of restriction fragments and mark the DNA ends, we incubated each sample with 37.5 μl of 0.4 mM biotin- 14-dCTP and 1.5 μl of 10 mM dATP, dGTP, and dTTP (all from Life Technologies) plus 40 units of Klenow fragment (NEB) at 23°C for 1.5 hours. We then performed ligation with 2,000 units of T4 DNA ligase (NEB) at 23°C for 4 hours with slow rotation. After the samples were treated with proteinase K (NEB) and 10% SDS, the chromatin was decrosslinked overnight in the presence of 250 mM NaCl at 68°C. DNA was purified using AMPure XP beads (Beckman Coulter) and sheared to 300-500 bp using a focused ultrasonicator (Covaris S220). The biotinylated DNA was selectively purified using Dynabeads MyOne Streptavidin T1 beads (Life Technologies). Hi-C library construction (end repair, adenylation, and adapter ligation) of the biotinylated DNA was done manually and adapter ligation was done using a TruSeq DNA Index kit (Illumina). The Hi-C library was quantified using a KAPA library quantification kit (Roche), and further PCR amplification was performed using Phusion Hot Start II DNA polymerase (Thermo Scientific). Finally, the library was sequenced with an Illumina HiSeq 2500 and HiSeq 4000 to obtain either 75-bp or 100-bp paired end reads, respectively.

The *in situ* Hi-C experiments for Control, *Exosc10KD*, and *Rad21KD* mESCs (1 million cells each) were performed according to the Arima-HiC protocol described in the Arima-HiC+ kit (Arima Genomics). The Illumina-compatible Hi-C libraries were prepared according to the Arima-HiC protocol and the library was sequenced with an Illumina Novaseq6000 using the pair-end method (150-bp reads).

### *In situ* Hi-C analysis in fission yeast

#### Basic Hi-C analysis

The sequenced paired-reads were analyzed using a HiC-Pro (Servant et al., 2015) workflow which is an integrated tool that can be used to preprocess Hi-C data by mapping, filtering, building the interaction matrix, and normalization. Statistics of the Hi-C data are shown in Supplemental Table 2. The raw paired-reads were mapped to the *S. pombe* reference genome (ASM294v2) using bowtie2 (Langmead and Salzberg, 2012) (v2.3.4). Hi-C artifacts (low mapping quality, multi-mapped reads, singletons, dangling end, self-circle, and dumped pairs) were removed and the valid read pairs between samples were then merged. To normalize the difference in sequencing depth from each Hi-C data, the *in situ* Hi-C data were normalized by the smallest number of valid read pairs. The raw and ICE-normalized matrix was generated at 1-kb and 3-kb bin resolutions using normalized valid pairs. To measure reproducibility among the replicates, we assessed the reproducibility either by comparing the contact probability using pearson correlation method or by measuring the correlation among Hi-C interaction matrices using a stratum-adjusted correlation coefficient (Yang et al., 2017).

#### Heatmap and plot

All heatmaps were generated based on the 3-kb iced-normalized matrix from HiC- Pro, using HiCPlotter (Akdemir and Chin, 2015) and HiCExplorer (Ramírez et al., 2018). The contact probability plot was calculated according to the genomic distance, and the quantitative comparisons of short-range (<100-kb) and long-range (>300-kb) contact probability plots were visualized using the ggplot2 package in R. For quantitative comparison of contact probability for each range (short-range and long-range), we used 1-kb resolution to obtain a sufficient number of bins for the analysis. Wilcoxon rank-sum tests were used to measure the statistical significance of each mutant compared to wild-type in terms of the short-range and long-range contact probabilities.

#### Potential border

To determine the potential border, we calculated the border strength (Kim et al., 2016; Van Bortle et al., 2014) for the *S. pombe* genome. First, we divided the *S. pombe* genome into 3-kb bins.

The range within ±5 bins (=15-kb) was set to ‘intra-domain’ for all bins in the genome, and the domain between upstream intra-domain and downstream intra-domain was termed as ‘inter-domain’. The score for each domain was calculated by the sum of the iced-normalized interaction score in the domain. The ratio between the two domains was calculated by dividing the intra-domain score by the inter-domain score. All bins were normalized using the z-score. The border strength score can explain the existence of a potential border; a positive strength score means that the interaction within the upstream and downstream intra-domain of a specific bin occurs more frequently than the interaction in the inter- domain.

#### Cohesin-mediated domain boundary

To generate cohesin-mediated meta border plots, the iced-normalized matrix was transformed into an obs/exp matrix (Vara et al., 2019). Then, the sub-matrix was extracted for the ± 50 kb region from the center of the convergent gene with strong Rad21 peaks. After all of the sub-matrices were averaged, the results were transformed with log10 and used to generate a meta plot via HiCExplorer.

### *In Situ* Hi-C analysis in mouse embryonic stem cells

#### Basic Hi-C analysis

Hi-C data were aligned to the mouse genome (mm10) and normalized using HiC- Pro. Statistics of the Hi-C data are shown in Supplemental Table 2. To achieve maximum sequencing depth, we pooled all replicates from the samples and proceeded to downstream analysis using R package GENOVA unless otherwise stated (https://github.com/robinweide/GENOVA) (Haarhuis et al., 2017).

#### Principal Component Analysis (PCA)

PCA was performed using HOMER (Heinz et al., 2018). Briefly, Hi-C paired-end reads were trimmed using HOMER (homerTools trim) and then separately aligned to the mouse genome (mm10) using bowtie2. Aligned Hi-C data were processed using HOMER (makeTagDirectory, runHiCpca.pl) with 100-kb resolution to achieve principal component 1 (PC1) values.

#### TAD calling

TAD calling was performed using CaTCH (Zhan et al., 2017). TADs were identified according to the reciprocal insulation (RI) value with the maximum CTCF enrichment at the TAD boundaries.

#### Contact domain and peak calling

Contact domain and peak calling were performed using JUICER (Durand et al., 2016). Contact domains with resolutions of 10, 20, 40, 50, and 100-kb were identified using the arrowhead algorithm (Rao et al., 2014). Peaks (chromatin loops) were identified using the HiCCUPS algorithm with default options (medium-resolution maps).

#### Aggregate TAD analysis (ATA)

ATA was performed using the R package GENOVA. Briefly, we used 20-kb resolution Hi-C matrices to calculate the average contact frequency matrices in the Control (si*Egfp*-treated mouse embryonic stem cell) TADs. To account for different TAD sizes, the Hi-C matrices were scaled to the same size, and the 5’ and 3’ TAD borders were extended to half of the TAD sizes. Finally, the differential ATA was produced by subtracting the average contact frequency matrices of knockdown (*Exosc10KD* or *Rad21KD*) and Control cells.

#### Aggregate peak analysis (APA)

APA was performed using the R package GENOVA. Briefly, we used 10-kb resolution Hi-C matrices to calculate the average contact frequency at the Control (si*Egfp*-treated mouse embryonic stem cell) peaks. The regions surrounding the Control peaks were extended as indicated in the figures.

#### Insulation score analysis

Insulation score analysis was performed using R package GENOVA. Briefly, genome-wide insulation scores (Crane et al., 2015) were calculated at 20-kb resolution with default options (window size = 25).

### Chromatin immunoprecipitation assay and ChIP-seq library preparation

ChIP assays (in fission yeast) were performed as previously described (Strahl-Bolsinger et al., 1997), with slight modifications. For the quantitative comparison of Rad21-5XFLAG ChIP in *rrp6*Δ and *wpl1*Δ background cells, we mixed formaldehyde-fixed *S. pombe* and *S. cerevisiae* at a 19: 1 ratio (OD600 ratio) before ChIP cell lysis step. Briefly, lysates were sonicated 12 times (15 sec on, 180 sec off) using a Dismembrator Model 500 sonicator (Fisher Scientific) at 30% output. The immunoprecipitated DNA was eluted using a SigmaPrep™ spin column and then subjected to protease treatment and decrosslinking. The extracted DNA was purified using a QIAquick PCR purification kit (Qiagen). The quantification of immunoprecipitated DNA relative to input was done by real-time ChIP-qPCR using Biorad CFX96 with EvaGreen Mix (BioFACT). All Primers used in the ChIP-qPCR were listed in Supplemental Table 4.

ChIP assays in mESCs were performed as previously described (Kim et al., 2014), with minor modifications. Briefly, mESCs were washed with PBS and incubated with 1% formaldehyde for 10 min at 25°C. The crosslinking was quenched with 125 mM glycine for 5 min at 25°C, and the cells were harvested with cold PBS. The cells were suspended in SDS lysis buffer (50 mM Tris-HCl, pH 8.0, 1% SDS, 10 mM EDTA) and sheared to mono- or dinucleosome sizes using. The sonicates were incubated overnight with the relevant antibodies and Protein A, G sepharose (17-1279-03 and 17-0618-05, GE Healthcare) at 4°C with agitation. The immune complexes were washed for 10 min each with the following wash buffers: low-salt wash buffer (20 mM Tris-HCl, pH 8.0, 150 mM NaCl, 0.1% SDS, 1% Triton X-100, 2 mM EDTA), high-salt wash buffer (20 mM Tris-HCl, pH 8.0, 500 mM NaCl, 0.1% SDS, 1% Triton X-100, 2 mM EDTA), and LiCl wash buffer (10 mM Tris-HCl, pH 8.0, 250 mM LiCl, 1% NP-40, 1% sodium deoxycholate, 1 mM EDTA). The immune complexes were further washed twice with TE buffer (10 mM Tris-HCl, pH 8.0, 1 mM EDTA), eluted with elution buffer (1% SDS, 0.1 M NaHCO3), and decrosslinked overnight at 68°C. The immunoprecipitated DNA was treated with proteinase K and RNase A and recovered by phenol-chloroform-isoamyl alcohol precipitation.

Sequencing libraries were prepared using either a NEXTflex™ Illumina ChIP-Seq Library Prep kit (5143-02, BIOO) or Accel-NGS 2S Plus DNA Library Kit (21024, SWIFT) according to the manufacturer’s protocols. Sequencing was performed on a HiSeq2500 using the single-end method (50- bp reads) or a Novaseq6000 using the pair-end method (150-bp reads).

### ChIP-seq data analysis

The sequenced reads were mapped to the *S. pombe* genome (ASM294v2) using NovoAlign (Novocraft Technologies) with default parameters. For pair-end reads, only the R1 reads were used for analysis. Bigwigs were created using Deeptools (Ramírez et al., 2016) bamCoverage, with bamfiles normalized to the number of reads per bin by the number of mapped reads in million (RPM). The bigwigs were normalized by mock (non-tagged control). Raw IP data or normalized data were visualized by integrative genome viewer (IGV). Plots and heatmaps of ChIP-seq data were generated using computeMatrix and plotProfile/Heatmap from Deeptools. For cohesin peak calling, MACS2 (Feng et al., 2012) callPeak was used. Adjacent and multiple Rad21 peaks were annotated to 3’-end of convergent genes. When multiple peaks were annotated to a 3’-end of convergent genes, a peak with maximum intensity at the converging intergenic sites was selected.

### Growth assay

Cells were spotted onto solid YES medium at 5-fold serial dilutions (in water) using a 48-pin replicator, and the plates were incubated at 30°C for 3 to 4 days.

### Protein extraction and Western blot analysis

To check the expression level of Rad21 by SDS-PAGE gel analysis, whole-cell protein extracts were prepared either by a standard extraction protocol (Matsuo et al., 2006) or by a bead-beating method in which cells were resuspended with Laemmli sample buffer (4% SDS, 40 mM Tris-HCl, pH 6.8, 5% glycerol, 5% *β*-mercaptoethanol, 0.04% bromophenol blue) and lysed with glass beads.

### Microscopic analysis

Prepared yeast colonies were spread on a coverglass-bottom dish (100350, SPL) and covered with a drop of soft agar. Images of whole yeast cells were captured by Z-stacking at a depth of 6 μm and steps of 0.3 μm, using a 488-nm laser and a Nikon A1R plus confocal microscope fitted with a CFI plan Apochromat λ DM 100x oil objective lens (1.45 numerical aperture, Nikon). Images were acquired at a resolution of 1024 x 1024 using a 3x digital zoom. GFP images were further evaluated with an Olympus IX-81 inverted fluorescence microscope.

### Real-time reverse transcrption PCR and library preparation for mRNA-sequencing

Total RNA was extracted using the hot-phenol method (Schmitt et al., 1990) and treated with recombinant DnaseI (Ambion). For real-time reverse transcription PCR, either an oligo(dT)15 primer or a gene-specific primer were used with ImPromII Reverse Transcriptase (Promega). All primers used in RT-qPCR were listed in Supplemental Table 4.

For RNA-seq, DNase-treated mRNA was purified using an NEBNext® Poly(A) mRNA Magnetic Isolation Module (E7490S, NEB) and subjected to library preparation using a NEXTflex™ Rapid Directional mRNA-Seq kit (5138-10, BIOO) according to the manufacturer’s instructions. Each library was sequenced on a HiSeq2500 using the single-end method (50-bp reads). The sequenced reads were aligned to the *S. pombe* genome (ASM294v2) using the STAR aligner (Dobin et al., 2013). Strand-specific signal tracks of RNA-seq were generated by using bam2wig.py from RSeQC (Wang et al., 2012) with options --wigsum = 1,000,000,000.

### Small RNA purification and library preparation for small RNA-sequencing

Small RNAs preparation and small RNA-seq were performed as previously described (Seo et al., 2017). Briefly, small RNAs were purified and subjected to library construction using a NEXTFlex small RNA- seq (5132-05, BIOO) according to the manufacturer’s instructions. Each library was sequenced on a HiSeq2500 using the single-end method (50-bp reads). The small RNA-seq reads were further analyzed with the same method adopted in mRNA-sequencing analysis.

### Antibodies

The following antibodies were used for the immunoblot or chromatin immunoprecipitation analyses: anti-β-actin (sc47778, Santa Cruz), anti-FLAG (F1804, Sigma), anti-H3K9me3 (ab8898, Abcam), anti- Rad21 (ab992, Abcam), anti-CTCF (07-729, Millipore), anti-V5 (MCA1360, Bio-Rad), and anti-CTD-Ser2P (ab5095, Abcam).

### Data and Software Availability

The RNA-seq, ChIP-seq, and Hi-C seq datasets have been deposited to the GEO under accession number GSE85147. Correspondence and materials requests should be addressed to D.L.

## Figure supplements

**Figure 1 – figure supplement 1.**
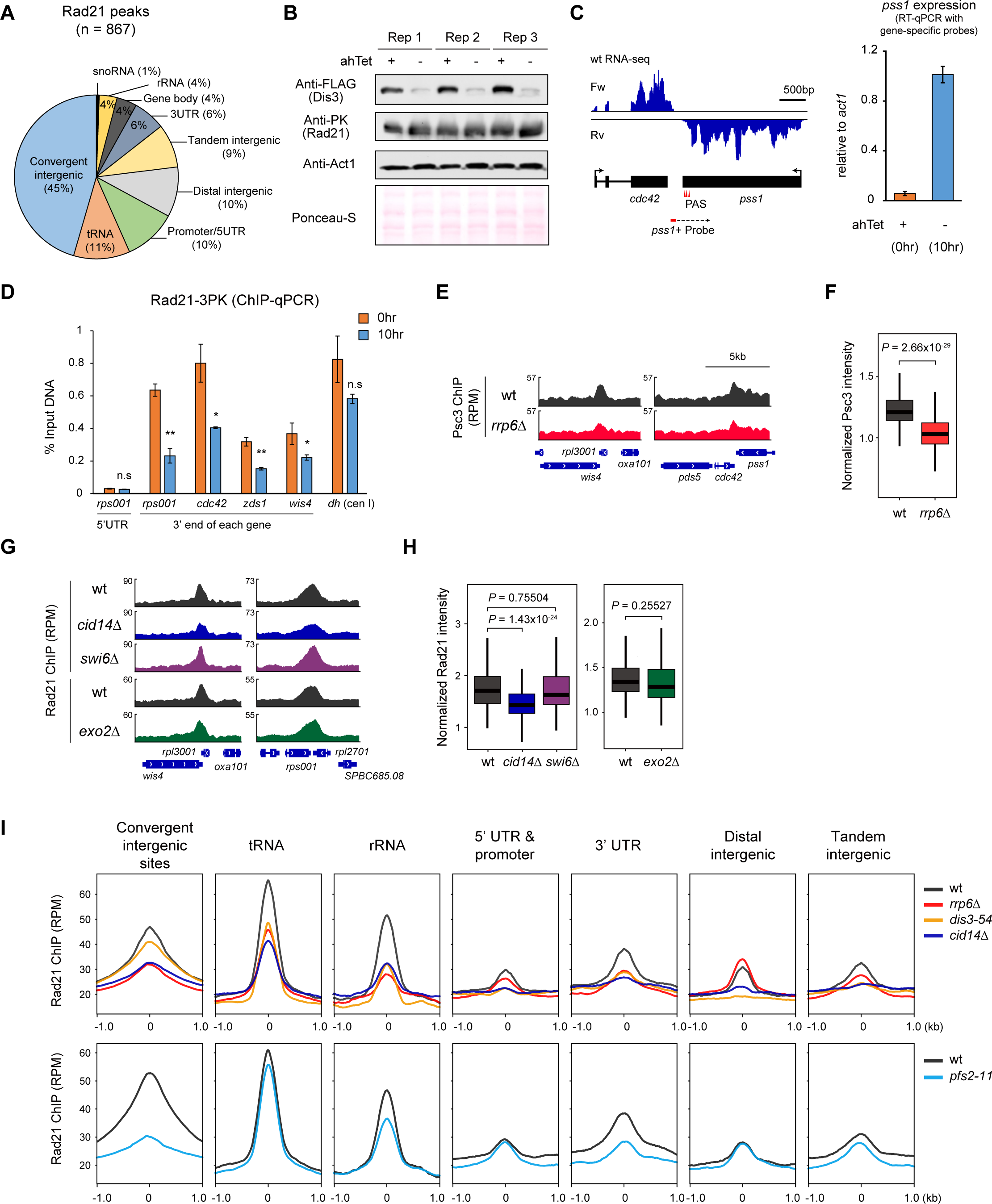
(A) Classification of Rad21 binding (n =867) sites at euchromatin. ‘Tandem intergenic’ indicates the intergenic regions between a pair of tandemly oriented genes. **(B)** Western blot validating the successful knockdown of Dis3 expression. Knockdown of Dis3 was achieved by incubating cells at rich medium (without anhydrotetracycline) for 10 hours. Ponceau S staining was used for the confirmation of equal loading. **(C)** RT-qPCR analysis showing the readthrough transcription at *pss1*locus by reverse- transcribing with a gene-specific primer (*pss1* probe) which targets downstream of Polyadenylation site in wt (0hr) and knockdown (10hr) of Dis3. **(D)** ChIP-qPCR analysis of Rad21 intensity upon knockdown (10hr) of Dis3. 5’UTR region of *rps001* was used as a negative control. Three biological replicates were used for (C) and (D). *P values* were calculated using the two-paired student t-test. **(E, F)** Representative examples (E) and box plot analysis (F) of Psc3 occupancy enriched at the 3’-end of convergent genes in wt and *rrp6*Δ**. (G, H)** Representative examples (G) and box plot analysis (H) of Rad21 occupancy enriched at the 3’-end of convergent genes in indicated strains. Each ChIP signal from box plot analysis (F, H) was normalized to that of the non-tagged control. **(I)** Metaplot analysis of Rad21 ChIP signals at indicated regions in various RNA processing mutants.

**Figure 1 – figure supplement 2.**
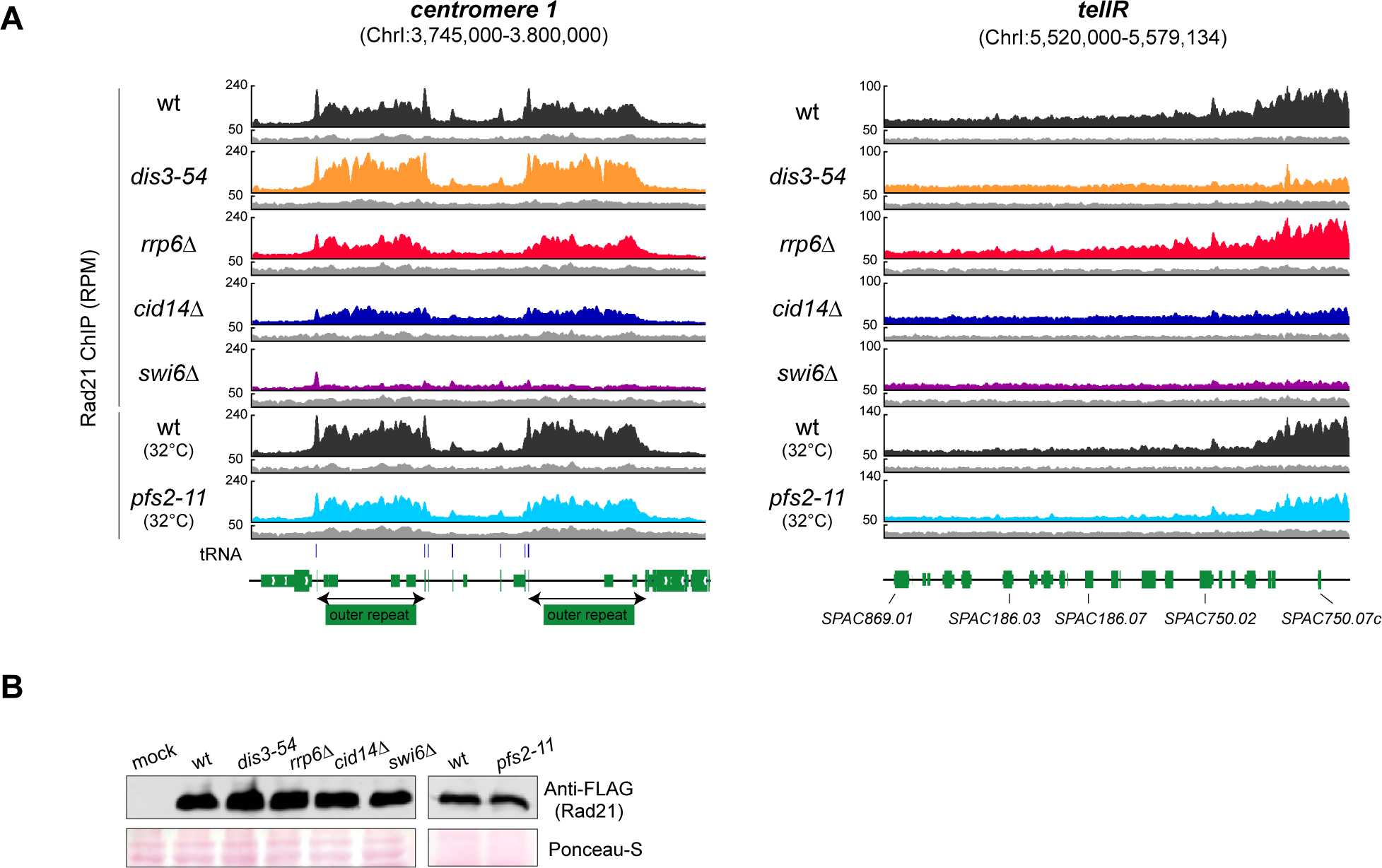
(A) Genome browser showing Rad21 ChIP-seq results at the centromere and telomere (right arm) of chromosome I. **(B)** Protein expression of FLAG-tagged Rad21 was unaffected in mutants, as assessed by Western blot analysis. Ponceau-S staining was used for the confirmation of equal loading.

**Figure 2 – figure supplement 1.**
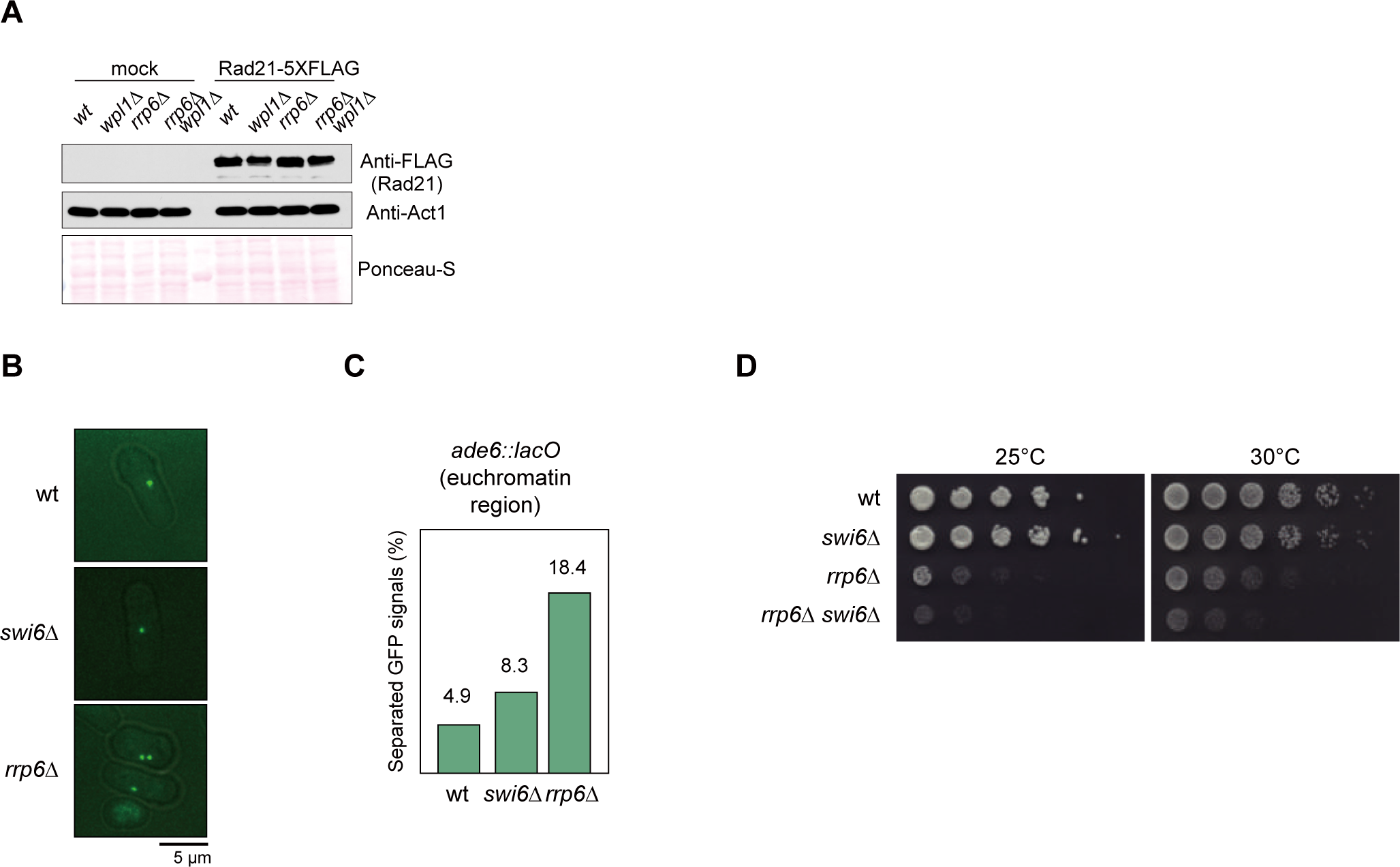
(A) Protein expression of FLAG-tagged Rad21 was unchanged in *rrp6Δ*, *wpl1Δ*, and *rrp6Δ wpl1Δ* when compared to wt. Ponceau-S staining was used for the confirmation of equal loading. **(B)** GFP spots indicating LacI-GFP bound to LacO repeats inserted at *ade6*^+^ site, which resides in chromosomal arm region. Merged GFP and DIC images are shown. (**C)** Bar plot shows the percentage of cells with two separate GFP foci from among more than 100 cells. GFP spots were counted from images captured at 0.3-μm intervals in the z-axis. **(D)** A spotting assay showing that Rrp6 genetically interacts with Swi6.

**Figure 3 – figure supplement 1.**
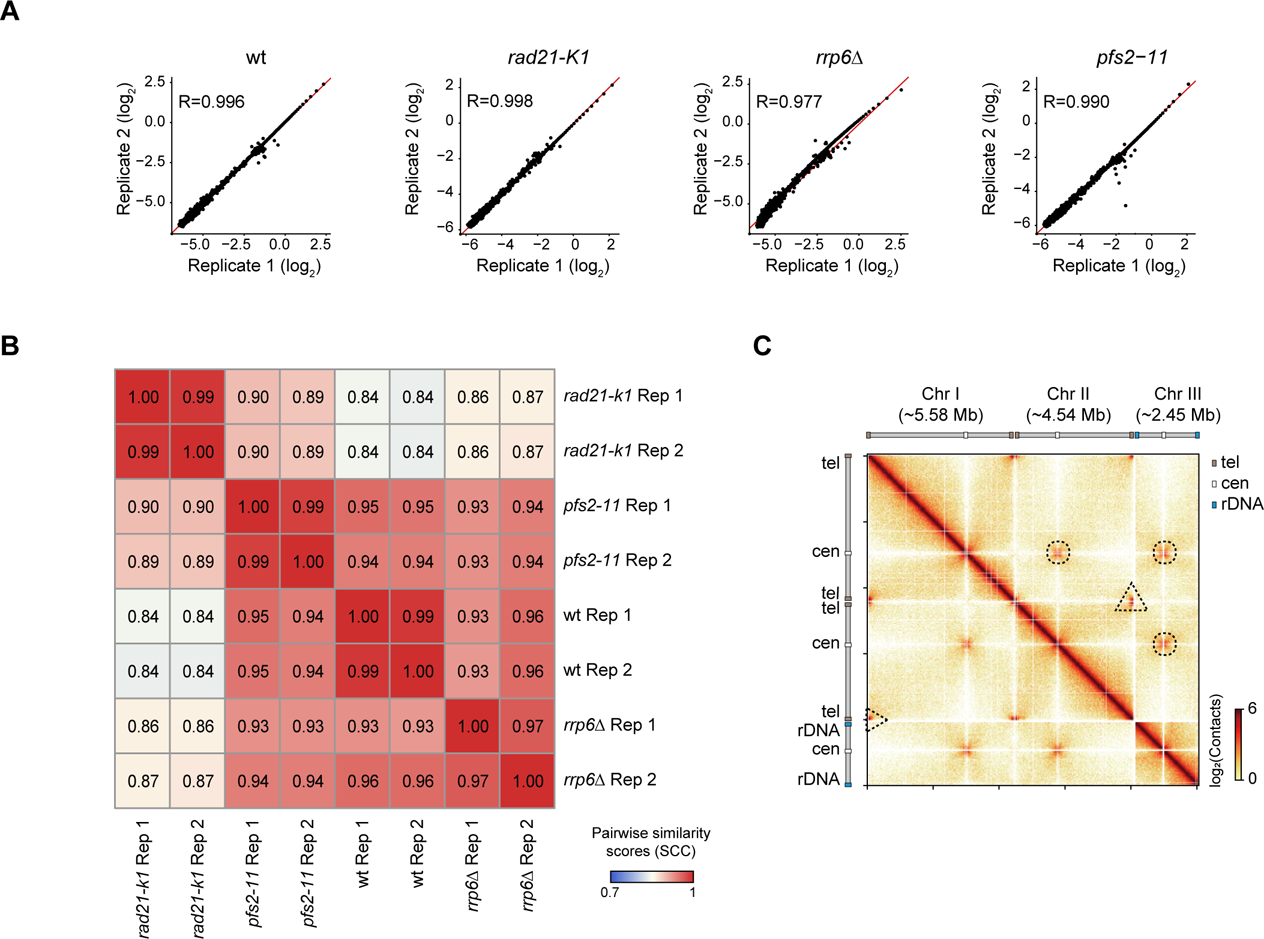
(A) Scatter plots showing high reproducibility among the replicates of all Hi-C samples, as assessed using contact probability based on Pearson correlation. (B) A stratum adjusted correlation coefficient (SCC) was used to compare the similiary across the replicate samples and each mutant using ICE- normalized Hi-C interaction matrices. **(C)** Genome-wide *in situ* Hi-C map of wild-type cells at 3-kb resolution showing the Rabl configuration. Dashed circles inside the Hi-C map indicate cen-cen and dashed triangles indicate tel-tel interactions.

**Figure 3 – figure supplement 2.**
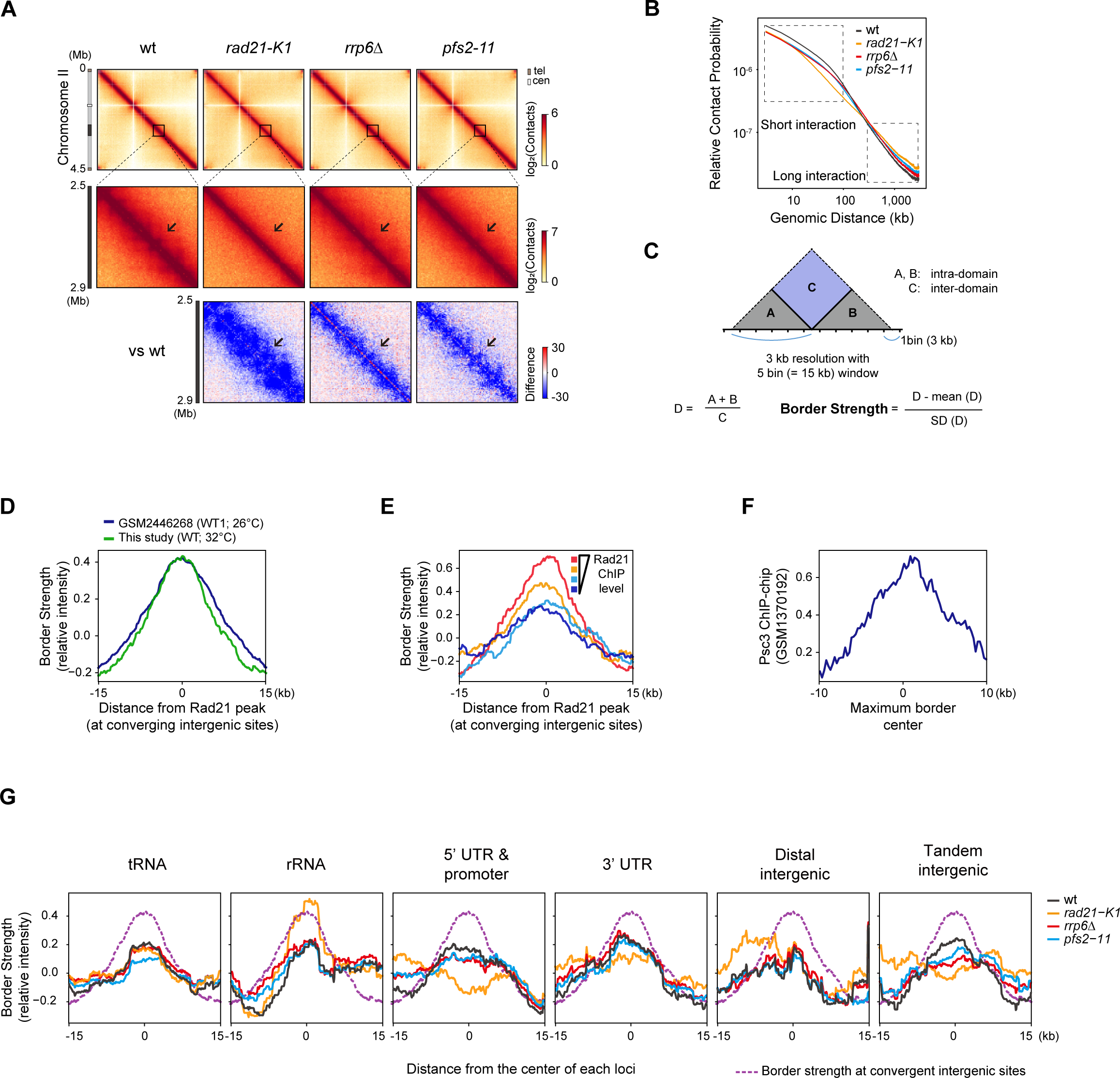
(A) *In situ* Hi-C map (3-kb resolution) in wt, *rrp6*Δ, *pfs2-11*, and *rad21-K1* showing the interaction in chromosome II (top) and the 2,500 - 2,900 kb region of chromosome II (bottom). **(B)** Relative contact probability (RCP) plot showing contact frequency relative to genomic distance. The dashed boxes display short-range (<100-kb) and long-range (>300-kb) interactions. **(C)** Scheme of border strength calculation (Van Bortle et al., 2014). **(D)** Enrichment of border strength calculated from this study (wt) and published data (GSM2446268) on Rad21 peaks that reside in converging intergenic sites. **(E)** Metaplot analysis of border strength according to the Rad21 peaks grouped by Rad21 ChIP intensity. **(F)** Enrichment of Psc3 ChIP-chip (GSM1370192) at the center of border in wt. **(G)** Metaplot analysis of the border strength at Rad21-enriched loci. Each plot was plotted against the center of Rad21 peaks overlapping with annotated regions. A purple dotted line indicates the border strength profile at convergent intergenic sites.

**Figure 3 – figure supplement 3.**
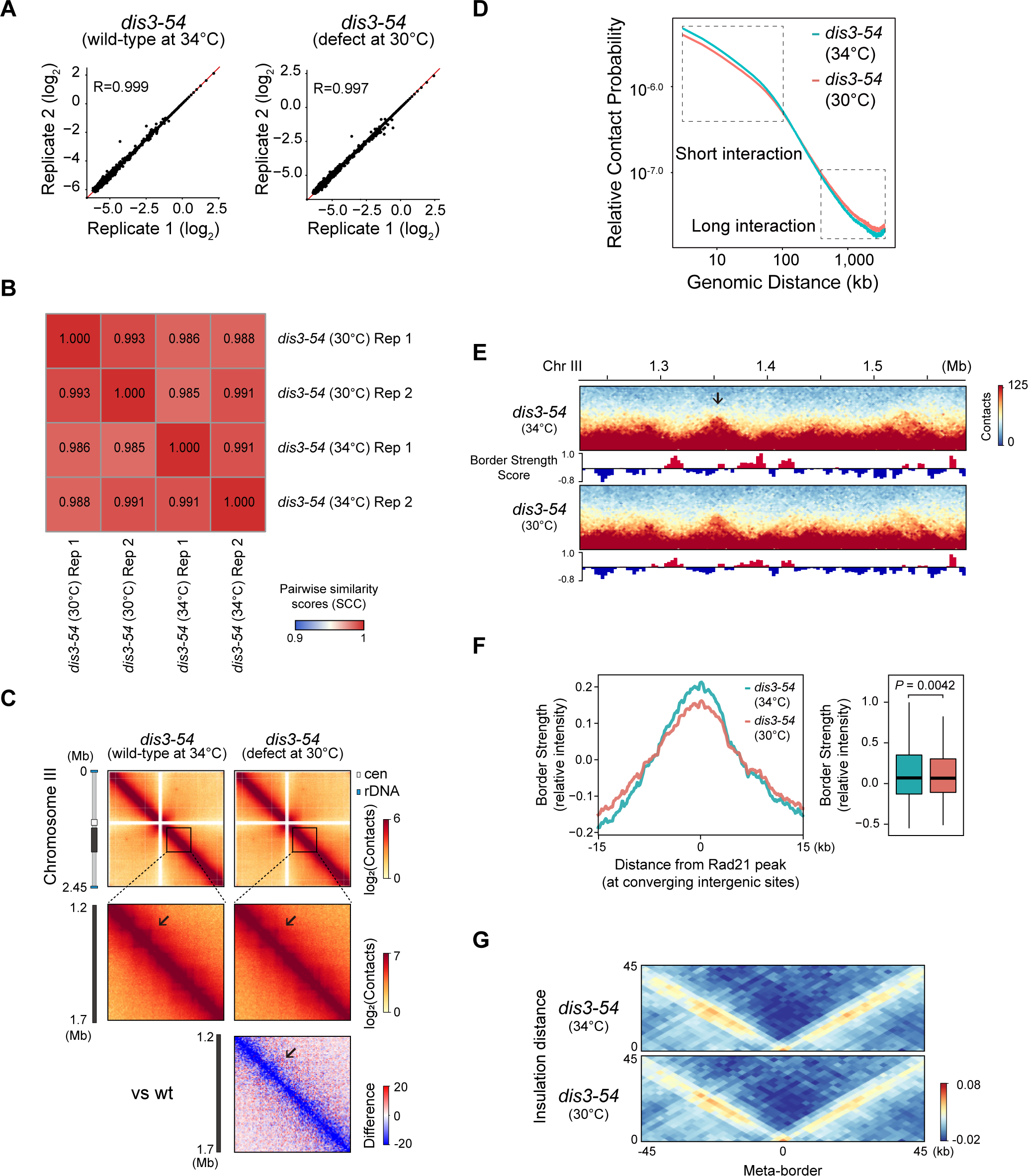
(A) Scatter plots showing high reproducibility in *dis3-54* cells using Pearson correlation analysis as in Figure 3 – figure supplement 1A. **(B)** A stratum adjusted correlation coefficient (SCC) was used to compare the similiary across the replicate samples and each mutant using ICE-normalized Hi-C interaction matrices. **(C)** *In situ* Hi-C map (3-kb resolution) in *dis3-54* mutants showing the interaction in chromosome III (top) and the 1,200 - 1,700 kb region of chromosome III (bottom). A relative Hi-C map comparing the mutants with that of wt was calculated by subtraction. **(D)** RCP plot showing contact frequency relative to genomic distance. The dashed boxes display short-range (<100-kb) and long-range (>300-kb) interactions. **(E)** Representative Hi-C map showing the 1,200 - 1,700 kb region of chromosome III (3-kb resolution). Border Strength is calculated with 3-kb resolution of 15-kb window sizes. Arrows indicate notable changes in the interaction domains. **(F)** Metaplot (Left) and boxplot (Right) of border strength at Rad21 peaks at converging intergenic sites **(G)** Aggregate plots generated from Hi-C data in *dis3-54* mutants at convergent gene pairs enriched with strong Rad21 peaks (n = 292) showing cohesin-mediated border boundary.

**Figure 4 – figure supplement 1.**
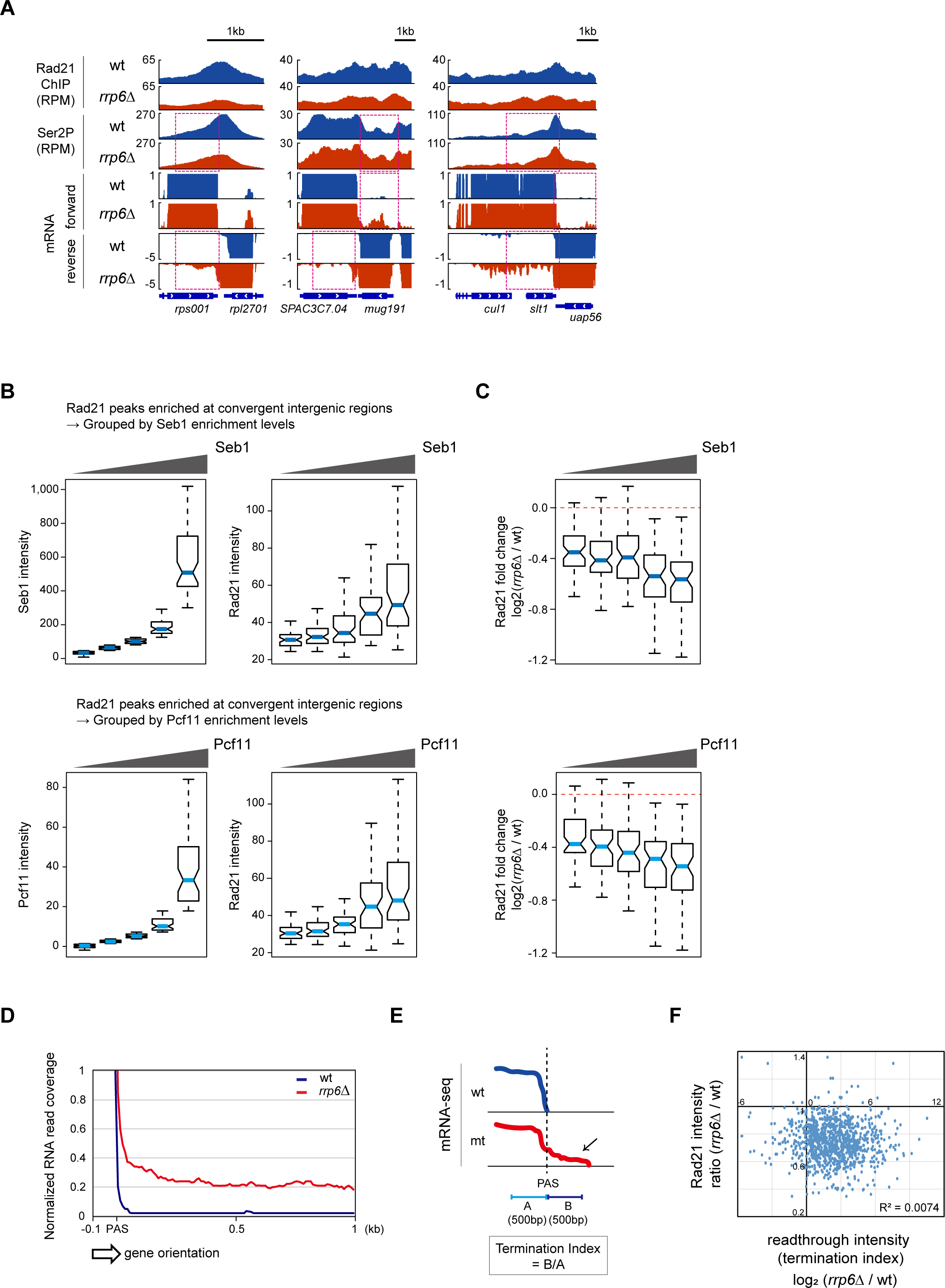
(A) Representative examples of genome browser views showing shifted Ser2P and decreased Rad21 ChIP intensity along with increased readthrough transcripts in *rrp6*Δ (dotted red box) **(B)** Comparison of Rad21 enrichment level according to five groups sorted by the level of Seb1 (A, top) and Pcf11 (A, bottom) at Rad21 peaks at convergent intergenic regions. **(C)** Log2 fold change of Rad21 (*rrp6*Δ/wt) were plotted by the aforementioned groups in (B). **(D)** Metaplot of mRNA-seq signals downstream of PAS against Rad21-enriched convergent genes. **(E)** To quantitatively compare termination defects (in terms of readthrough transcript) in *rrp6*Δ with wt, we used termination index to indicate the readthrough intensity. Termination index (TI) is determined by dividing B (the read count within PAS + 500 bp) by A (the read count within PAS - 500bp). PAS, Polyadenylation signal. **(F)** Scatter plot comparing the change in readthrough intensity (calculated as the termination index) as log2 (*rrp6*Δ/wt) (x-axis) with that of Rad21 occupancy as *rrp6*Δ/wt (y-axis).

**Figure 5 – figure supplement 1.**
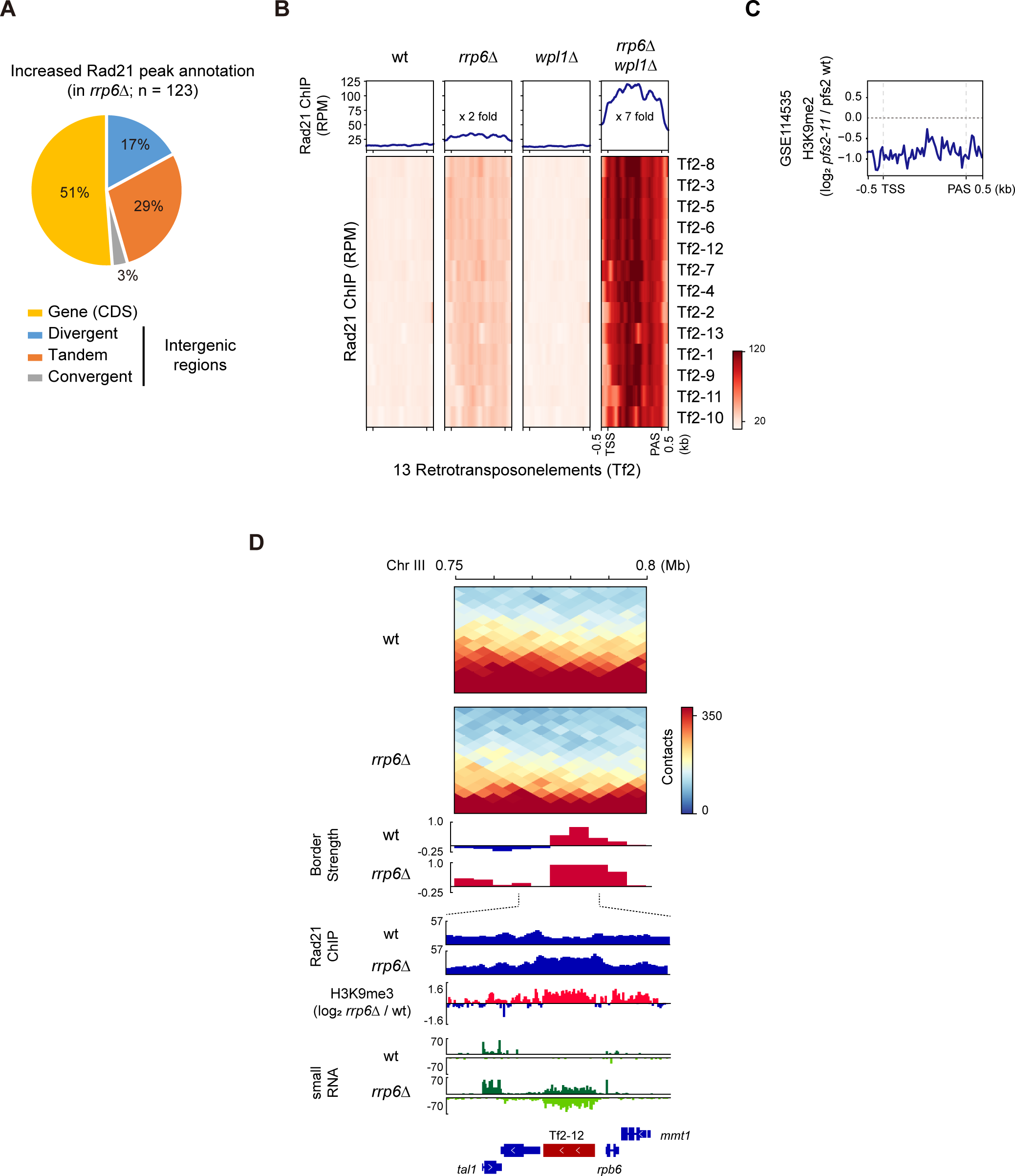
(A) Annotation of increased Rad21 peaks upon Rrp6 deletion. **(B)** Heatmaps of Rad21 ChIP-seq signals (RPM) in indicated strains at 13 retrotransposon elements. Relative fold change of Rad21 signals in *rrp6*Δ and *rrp6*Δ *wpl1*Δ compare to wt was compared by calculating median Rad21 intensity at 13 Tf2 ORFs. **(C)** Metaplot analysis comparing H3K9me2 levels in *pfs2-11* mutant (2hr heat shock at 37°C) and wt state. The H3K9me2 data were derived from the published dataset (GSE114535). **(D)** *In situ* Hi- C map (5-kb resolution) (top), the border strength (middle), and chromatin features (bottom) at retrotransposon element *Tf2-12* at wt and *rrp6*Δ.

**Figure 6 – figure supplement 1.**
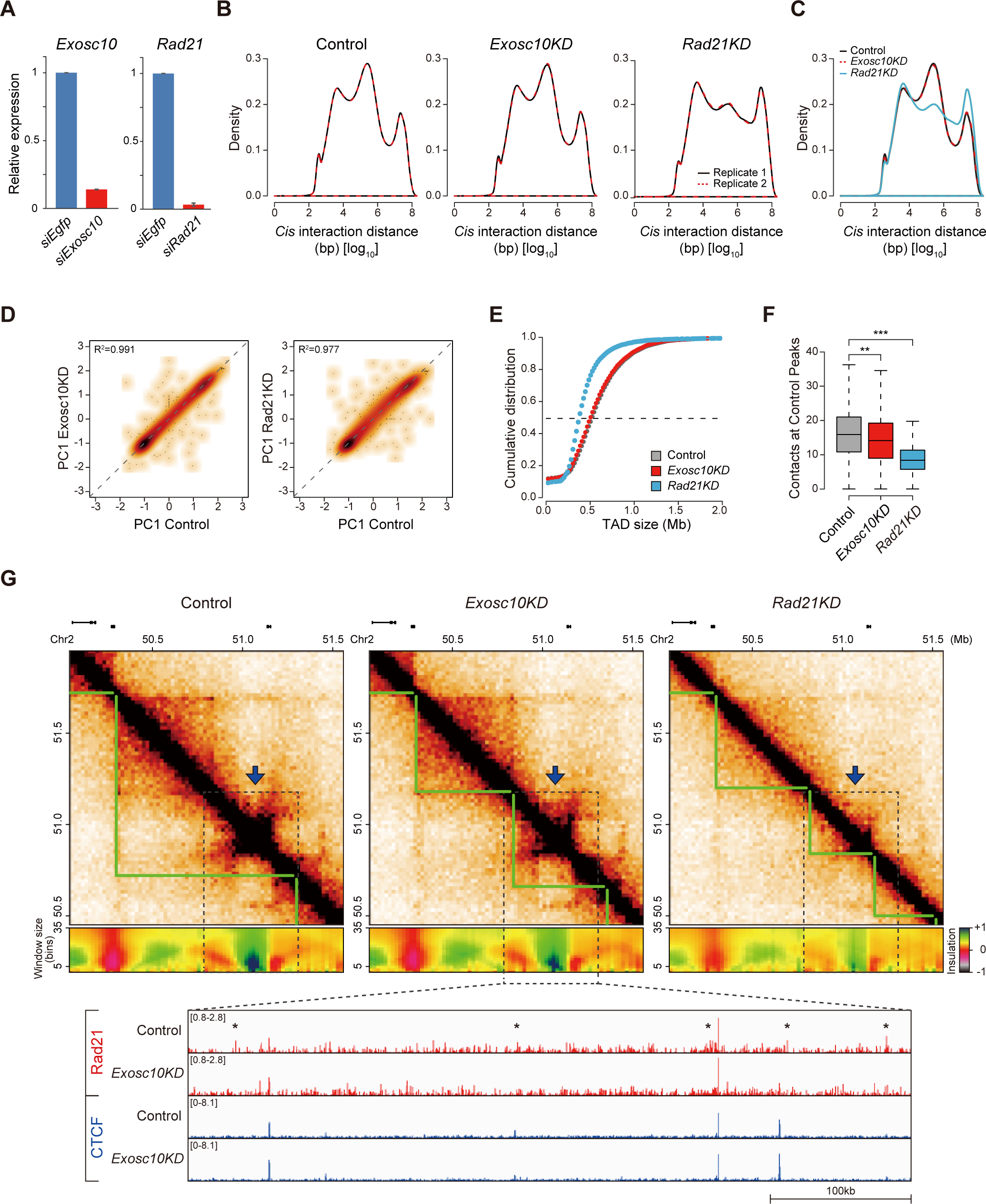
(A) Bar plot showing the knockdown efficiencies of siRNA against *Exosc10* and *Rad21*. The levels of the target genes (*Exosc10* and *Rad21*) were normalized relative to that of *Act1*. Data are presented as mean ± SD. **(B)** Density plots showing the frequency of *cis* interactions of two replicates in Control (left), *Exosc10KD* (middle), and *Rad21KD* (right) mESCs. Black and red dotted lines indicate replicates 1 and 2, respectively. **(C)** Density plot showing the frequency of cis interactions of Control (black line), *Exosc10KD* (red dotted line), and *Rad21KD* (light blue line) mESCs. **(D)** Scatter plots comparing genome-wide PC1 (first principal components, equivalent to the first eigenvectors) values between *Exosc10KD* (left, Pearson R^2^=0.991) and *Rad21KD* (right, Pearson R^2^=0.977) with Control cells. **(E)** Cumulative distribution of TAD sizes in Control (gray dots), *Exosc10KD* (red dots), and *Rad21KD* (light blue dots) mESCs. The dashed line indicates average TAD size. **(F)** Box plot displaying contacts (interactions) of Control (gray), *Exosc10KD* (red), and *Rad21KD* (light blue) mESCs at Control peaks (chromatin loops). The horizontal line in the box denotes the median. *P* values were calculated using the paired Wilcoxon rank-sum test (***P < 10^-100^; **P < 10^-50^). **(G)** Example of a TAD that lost its interactions (dashed box with blue arrow) upon Exosc10 and Rad21 knockdown. Hi-C contact maps (top), insulation heatmaps (middle), and Rad21 and CTCF ChIP-seq peaks (bottom, red and blue bars, respectively) are shown for Control, *Exosc10KD*, and *Rad21KD* ESCs. The ChIP-seq tracks display zoom-in browser view of the dashed box, showing where a TAD loses its interactions. The decreased Rad21 ChIP-seq peaks upon *Exosc10KD* are marked with asterisks.

