## Supplemental Information for "RNA surveillance controls 3D genome structure via stable cohesin-chromosome interaction"

### Supplemental Table 1. List of yeast strains used in this study

| Strain | mating type | Genotype | Source |
|---|---|---|---|
| 972 | *h-* | 972 | |
| ED665 | *h-* | *ade6-M210 leu1-32 ura4-D18* | This study |
| #4277 | *h-* | *rrp6Δ::kanMX6 ade6-M210 leu1-32 ura4-D18* | This study |
| #2981 | *h-* | *rrp6Δ::kanMX6 Rad21-5XFLAG::hphMX6 ade6-M210 leu1-32 ura4-D18* | This study |
| #5632 | *h-* | *rrp6Δ::kanMX6* | This study |
| #3104 | *h-* | *rad21-K1:hygR #19* | Mitsuhiro Yanagida |
| #2709 | *h-* | *Rad21-5XFLAG::hphMX6 ade6-M210 leu1-32 ura4-D18* | This study |
| #3670 | *h-* | *pfs2-11 Rad21-5XFLAG::hphMX6* | This study |
| HU2644 | *h-* | *pfs2-11 (HU2644)* | Karl Ekwall |
| RC6519 | *h-* | *dis3 leu1-32?=ALP344* | Robin Allshire |
| #3373 | *h-* | *dis3 leu1-32?=ALP344 x Rad21-5XFlag:: hphMX6* | This study |
| FY9143[a] | h- | *leu1 dis3-54* | NBRP |
| #3918 | *h+* | *lys1-131 ura4-D18 ade6[::kanr-ura4+-lacOp] his7+::lacI-GFP rrp6Δ:: hphMX6* | This study |
| #3590 | *h+* | *lys1-131 ura4-D18 ade6[::kanr-ura4+-lacOp] his7+::lacI-GFP swi6Δ:: hphMX6* | This study |
| FY15598[a] | *h+* | *lys1-131 ura4-D18 ade6[::kanr-ura4+-lacOp] his7+::lacI-GFP* | NBRP |
| #1090 | *h-* | *swi6Δ::kanMX6 ade6-M210 leu1-32 ura4-D18* | This study |
| #2738 | *h-* | *swi6Δ::kanMX6 Rad21-5XFLAG::hphMX6 ade6-M210 leu1-32 ura4-D18* | This study |
| #3379 | *h+* | *rrp6Δ::hphMX6 swi6Δ:: kanMX6 ade6-M210 leu1-32 ura4-D18* | This study |
| #4069 | *h-* | *Psc3-5XFlag::kanMX6 ade6-M210 leu1-32 ura4-D18* | This study |
| #4071 | *h-* | *rrp6Δ::hphMX6 Psc3-5XFlag::kanMX6 ade6-M210 leu1-32 ura4-D18* | This study |
| #4079 | *h-* | *cid14Δ::LEU2MX6 leu1-32 ura4-D18 Rad21-5XFlag::hphMX6* | This study |
| #4097 | *h-* | *exo2Δ::kanMX6 Rad21-5XFlag::hphMX6 ade6-M210 leu1-32 ura4-D18* | This study |
| #5886 | *h-* | *tet*$O_7$TATA$_{cyc1}$-3XFLAG-*Rrp6::hphMX6* pDM291-*tet*R-*tup11Δ70::*$ura4^+$ | This study |
| #6121 | *h-* | *tet*$O_7$TATA$_{cyc1}$-3XFLAG-*Rrp6::hphMX6 Rad21-3XPK::natMX6* pDM291-*tet*R-*tup11Δ70::*$ura4^+$ | This study |
| #6125 | *h-* | *tet*$O_7$TATA$_{cyc1}$-3XFLAG-*Dis3::hphMX6* pDM291-*tet*R-*tup11Δ70::*$ura4^+$ | This study |
| #6317 | *h-* | *tet*$O_7$TATA$_{cyc1}$-3XFLAG-*Dis3::hphMX6 Rad21-3XPK::natMX6* pDM291-*tet*R-*tup11Δ70::*$ura4^+$ | This study |
| #5965 | *h-* | *wpl1Δ::hphMX6 ade6-M210 leu1-32 ura4-D18* | This study |
| #6114 | *h-* | *rrp6Δ::hphMX6 ade6-M210 leu1-32 ura4-D18* | This study |
| #6234 | *h-* | *rrp6Δ::kanMX6 wpl1Δ::hphMX6 ade6-M210 leu1-32 ura4-D18* | This study |
| #6189 | *h-* | *rrp6Δ::natMX6 Rad21-5XFLAG::hphMX6 ade6-M210 leu1-32 ura4-D18* | This study |
| #6192 | *h-* | *wpl1Δ::kanMX6 Rad21-5XFLAG::hphMX6 ade6-M210 leu1-32 ura4-D18* | This study |
| #6203 | *h-* | *rrp6Δ::natMX6 wpl1Δ::kanMX6 Rad21-5XFLAG::hphMX6 ade6-M210 leu1-32 ura4-D18* | This study |
| #4770 | *h-* | *tfs1Δ::natMX6 ade6-M210 leu1-32 ura4-D18* | This study |
| #4772 | *h-* | *tfs1Δ::natMX6 Rad21-5XFLAG::hphMX6 ade6-M210 leu1-32 ura4-D18* | This study |

[a] The strain was purchased from the Yeast Genetic Resource Center of Japan supported by the National BioResource Project (YGRC/NBRP) (http://yeast.lab.nig.ac.jp/nig.v2.1)

**Supplemental Table 2.** ***in situ*** **Hi-C library information**

| Type | Valid interaction | *Cis* long-range (>5kb) | *Cis* short-range (≤5kb) | *Trans* interaction |
|---|---|---|---|---|
| wt | 42,985,517 | 25,625,958 | 10,102,783 | 7,256,776 |
| *rrp6Δ* | 33,561,228 | 19,338,964 | 6,873,478 | 7,348,786 |
| *rad21-K1* | 43,029,057 | 23,288,347 | 8,258,770 | 11,481,940 |
| *pfs2-11* | 48,019,476 | 27,979,882 | 9,343,382 | 10,696,212 |
| *dis3-54* (34°C) | 56,563,055 | 31,569,208 | 12,825,509 | 12,168,338 |
| *dis3-54* (30°C) | 50,561,641 | 28,314,922 | 9,747,823 | 12,498,896 |

| Type | Valid interaction | *Cis* long-range (>20kb) | *Cis* short-range (≤20kb) | *Trans* interaction |
|---|---|---|---|---|
| Control | 557,091,350 | 328,632,523 | 158,521,582 | 69,937,245 |
| *Exosc10KD* | 528,762,027 | 308,727,097 | 149,682,448 | 70,352,482 |
| *Rad21KD* | 528,504,739 | 307,326,002 | 150,680,921 | 70,497,816 |

Valid read-pairs were processed from HiC-Pro. The *in situ* Hi-C experiments were performed in biological duplicates and aligned replicates were then merged to generate a genome-wide interaction map.

**Supplemental Table 3. Summary of contact domain and chromatin loop identification in mESC *in situ* Hi-C**

| Contact Domains | Control | *Exosc10KD* | *Rad21KD* |
|---|---|---|---|
| 10 kb Resolution | 4,921 | 4,866 | 2,385 |
| 20 kb Resolution | 4,245 | 4,135 | 2,678 |
| 40 kb Resolution | 2,506 | 2,458 | 1,914 |
| 100 kb Resolution | 943 | 901 | 812 |
| 500 kb Resolution | 76 | 72 | 63 |
| **Loop** | **Control** | ***Exosc10KD*** | ***Rad21KD*** |
| Loops / peaks | 12,477 | 12,490 | 6,242 |

Contain domains were identified by Juicer arrowhead algorithm and chromatin loops (peaks) were identified by Juicer HiCCUPS algorithm.

### Supplemental Table 4. List of primers used in this study

| Targets | Sequences (5' - 3') | Purpose |
|---|---|---|
| sp_act1_rt_F | CCACTATGTATCCCGGTATTGC | RT-qPCR |
| sp_act1_rt_R | GAATGGATCCACCAATCCAGAC | RT-qPCR |
| sp_pss1-gene_specific_RT_R | AGGTGATACGATTATGTTGACT | RT-qPCR |
| sp_pss1_rt_F | CCTACCGAAACCCAGTTGAG | RT-qPCR |
| sp_pss1_rt_R | GGAAATTGACGAGCAAAAGG | RT-qPCR |
| sp_rps001_5UTR_F | TCCGCATGGATAACTAC | ChIP-qPCR |
| sp_rps001_5UTR_R | GTTGAAACAACGCAAAC | ChIP-qPCR |
| sp_rps001_3UTR_F | GCT GAG GCT CAA TAA ATG | ChIP-qPCR |
| sp_rps001_3UTR_R | GAC GGA GTT ACT CTA CTA AT | ChIP-qPCR |
| sp_cdc42_3UTR_F | GTATGGTCCGACTTTGA | ChIP-qPCR |
| sp_cdc42_3UTR_R | CGACAAGAGCAAATGAG | ChIP-qPCR |
| sp_zds1_3UTR_F | GGGAGTACGAAGTTACCCATTC | ChIP-qPCR |
| sp_zds1_3UTR_R | AGCACTTCAGTAGCGTTCTATG | ChIP-qPCR |
| sp_wis4_3UTR_F | AGAAGTGGATGTTGAGTGCAT | ChIP-qPCR |
| sp_wis4_3UTR_R | GCATGCAGATGATGTATTTACC | ChIP-qPCR |
| sp_cenI_dh_F | GTCAGCTCACTCAAGTCCAATC | ChIP-qPCR |
| sp_cenI_dh_R | CACTACAAGGACTAAGCCCAAG | ChIP-qPCR |
| mm_beta-actin_F | GGCTGTATTCCCCTCCATCG | RT-qPCR |
| mm_beta-actin_R | CCAGTTGGTAACAATGCCATGT | RT-qPCR |
| mm_Rad21_F | AGAGCATCATCTCACCAAAGG | RT-qPCR |
| mm_Rad21_R | GCCGAAACGCCATCTTTATTT | RT-qPCR |
| mm_Exosc10_F | GAGACACAAGGAGACAGGTTAC | RT-qPCR |
| mm_Exosc10_R | CATGCCCACTCTCTCCAATATC | RT-qPCR |
